# Quantum Spin Resonance in Engineered Magneto-Sensitive Fluorescent Proteins Enables Multi-Modal Sensing in Living Cells

**DOI:** 10.1101/2024.11.25.625143

**Authors:** Gabriel Abrahams, Ana Štuhec, Vincent Spreng, Robin Henry, Idris Kempf, Jessica James, Kirill Sechkar, Scott Stacey, Vicente Trelles-Fernandez, Lewis M. Antill, Christiane R. Timmel, Jack J. Miller, Maria Ingaramo, Andrew York, Jean-Philippe Tetienne, Harrison Steel

## Abstract

Quantum mechanical phenomena have been identified as fundamentally significant to an increasing number of biological processes. Simultaneously, quantum sensing is emerging as a cutting-edge technology for diverse applications across materials and biological science. However, until recently, biological based candidates for quantum sensors have been limited to *in vitro* systems, were prone to light induced degradation, and the experimental setups involved are typically not amenable to high-throughput study as would enable further engineering e.g. via directed evolution. We recently created a new class of magneto-sensitive fluorescent proteins (MFPs), which we show overcome these challenges and represent a new form of engineered biological quantum sensors that function both at physiological conditions and in living cells. Through directed evolution, we demonstrate the possibility of engineering these proteins to alter properties of their response to magnetic fields and radio frequencies. These effects are explained in terms of the radical pair mechanism (RPM), involving the protein backbone and a bound flavin cofactor. Using this engineered system we demonstrate the first observation of a fluorescent protein exhibiting Optically Detected Magnetic Resonance (ODMR) in living bacterial cells at room temperature, at sufficiently high signal-to-noise to be detected in a single cell. These magnetic resonance and magnetic field effects measured via fluorescence enable novel technologies; examples we demonstrate include spatial localisation of fluorescence signals using gradient fields (i.e. Magnetic Resonance Imaging (MRI) using a genetically encoded probe), sensing of the molecular microenvironment, multiplexing of bio-imaging, and lock-in detection, overcoming typical fluorescence imaging challenges of light scattering and autofluorescence. Taken together, our results represent a new range of sensing modalities for engineered biological systems, based on and designed around understanding the quantum mechanical properties of MFPs.

## 1 Introduction

Coupling electromagnetic fields with biological processes through fluorescence has revolutionised quantitative biology [1, 2]. Meanwhile, quantum sensing tools (i.e. those whose function arises from spin-dependent processes) have been developed for their unique advantages in biological applications, but have until recently been limited to realisations using non-biological probes [3–5] or measurements under *ex-vivo* conditions [6–8]. We previously reported the development of a library of magneto-responsive fluorescent protein (MFP) variants derived from the LOV2 domain (broadly termed MagLOV), which exhibit fluorescence signals with large magnetic field effects (MFEs) [Fig. 1 A] [9]. With this work, we demonstrate that at room temperature, we can detect optically detected magnetic resonance (ODMR) [10–19] in living cells expressing these fluorescent proteins, including at single cell level. The ODMR signature implies a quantum system whose properties and dynamics sensitively depend on the local environment, opening up a broad range of novel possibilities for cellular biosensing [3]. The ODMR arises from electron spin resonance (ESR) that we hypothesise originates from a spin-correlated radical pair (SCRP) [20, 21] [Fig. 1 B] involving the LOV2 domain non-covalently bound flavin cofactor chromophore. This theory is based on prior evidence [22] and supported by the MFE, ODMR, and spectral studies we present.

**Fig. 1.**
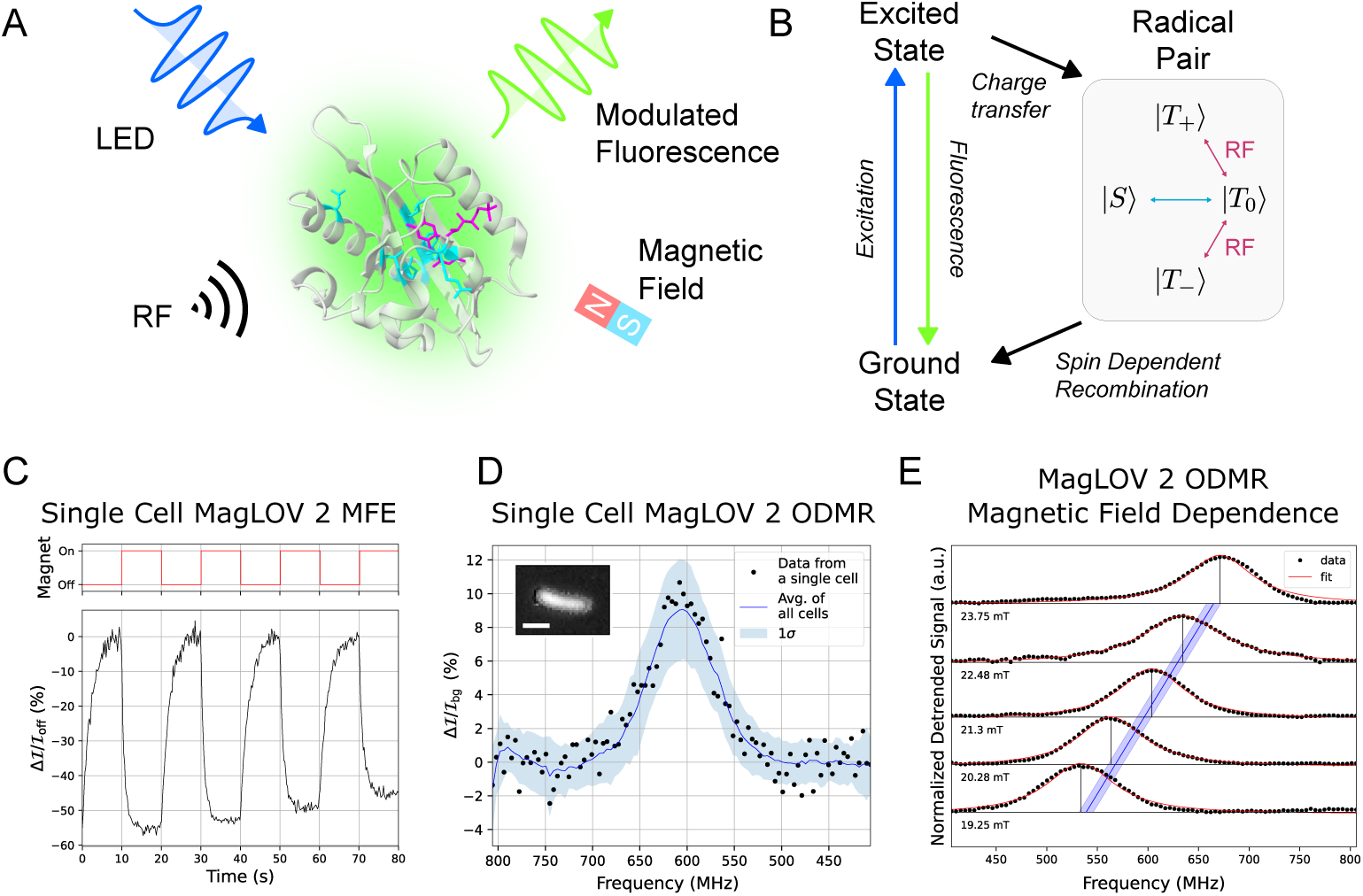
Observation of MFE and ODMR in magneto-sensitive fluorescent proteins. **A.** Structure of AsLOV2 PDB 2V1A[23] with mutations resulting in MagLOV 2 highlighted. Spin transitions driven by radio-frequency (RF) fields in the presence of a static magnetic field are optically detected via fluorescence measurements on an otherwise standard widefield microscope. **B.** Simplified photocycle diagram in the case of a large external magnetic field. The radical pair is born in a triplet state |*T* ⟩ with spin projections |*T*_0_⟩, |*T*_+_⟩ and |*T*_−_⟩[24, 25] and undergoes field-dependent singlet-triplet interconversion to the singlet state, |*S*⟩, driven by nuclear-electron spin-spin interactions. **C.** A single cell expressing MagLOV 2 displaying an MFE of ~ 50% (measured as a change in fluorescence intensity ℐ in the presence of an applied field). For MFE measurements, the magnetic field was switched between 0 mT and 10 mT, here with a period of 20 s. The intensity over time is integrated over pixels covering the single cell, with a background photobleaching trendline removed. Contrast is calculated as (ℐ − ℐ_off_)/ℐ_off_ = Δℐ/ℐ_off_. **D.** Black dots: data from a single cell expressing MagLOV 2 displaying an ODMR signal with ~ 10 % contrast. The static field *B*_0_ is ~21.6 mT. Blue line (shade): the mean (std) of all single cell data in a field of view (~1000 cells). Inset: microscope image cropped to a single cell expressing MagLOV 2 with a 2 µm scalebar. **E.** The static magnetic field *B*_0_ was varied by adjusting the magnet’s position, and ODMR spectra recorded. The red-lines are Lorentzian fits. The blue line is a theoretical prediction (i.e. is not a fit) of the expected resonance frequency of an electron spin with 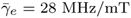 or *g_e_* = 2.00 with the shaded region representing uncertainty in the magnetic field strength (as determined by a Hall probe) at the sample position.

Both MFE and ODMR signals are straightforward to detect in cells on a standard wide-field fluorescence microscope, supporting further development and application of this discovery. Beyond the ease of detecting MFP’s magnetic resonance via emission, MFPs are advantageous over other candidate spin sensors for biological uses because they can be expressed directly in the host organism (e.g. allowing direct coupling and regulation by biological processes) and because their performance can be engineered genetically, such as through rational design or directed evolution. This engineerability is demonstrated through a selection approach previously reported [9] and here used to generate novel protein variants specialised for sensing applications.

We demonstrate applications of MFPs as reporters that can be used for lock-in signal amplification in noisy measurement environments (as often encountered in biological applications [26]), and to enable signal multiplexing by engineering variants with differing dynamic responses. We also show that the MagLOV MFE is attenuated by interaction with Magnetic Resonance Imaging (MRI) contrast agents, with a dose-dependent effect consistent with spin relaxation, implying MagLOV’s ability to sense the surrounding environment. Finally, we realise applications to spatial imaging; because the ODMR resonance condition depends on the static magnetic field at the location of the protein, it is possible to use gradient fields (as in MRI) to determine the spatial distribution of MFPs with scattering-independent measurements of a sample’s fluorescence. We demonstrate this by building a novel fluorescence-MRI instrument based on a small-animal MRI coil with a 1-dimensional magnetic gradient, which we apply to simultaneously localise the depth position of multiple bands of bacterial cells embedded in a ~40 cm^3^ volume.

The future potential for optimisation and application of MFPs is vast. For example, through high-throughput screening it is feasible to perform selection based on magnetic response, ODMR response, or biological factors such as protein stability. Down the line, the possibility of controlling charge state transfers within a protein raises the potential for developing 3D spatially-localised magneto-genetic actuators. Ultimately, the development of these systems may lead to a new paradigm of quantum-based tools for biological sensing, measurement, and actuation.

## 2 Optically Detected Magnetic and Resonant Field Effects in Fluorescent Living Cells

Reaction yield detected magnetic resonance, RYDMR (a form of ODMR) is both a diagnostic test for the existence of the proposed spin-correlated radical pair (discussed below) [27] and, due to its relative simplicity, an effective measurement modality for performing readout from quantum sensing devices in biological and materials applications [3, 4, 28–32]. ODMR studies the spin dynamics of such systems through the application of oscillating magnetic fields, *B*_1_, resonant with spin transition energies, and facilitates optical readouts through emission or absorption. Here we employ this technique to study the coherent spin dynamics of light-generated radical pairs whose photocycle contains a field-sensitive fluorophore, namely the flavin FMN.

Previous studies have reported MFEs arising from interactions between a flavin cofactor and protein in terms of the SCRP mechanism [33–41], with the most prominent example being cryptochromes implicated in avian magnetoreception [42–45]. Furthermore, radical pair intermediates are known to form in the LOV2 domain *Avena sativa* phototropin 1 (AsLOV2) variant C_450_A (the precursor protein to MagLOV) [46], as well as in related LOV domains [47–49]. Parallel with our results, ODMR has recently been reported in purified EYFP at room temperature, and in mammalian cells at cryogenic temperature [50]. Furthermore, subsequent preprints have reported ODMR in purified protein solutions of *Dm*Cry, mScarlet-flavin, and MagLOV at room temperature, as well as mScarlet-flavin in nematodes at room temperature [45, 51].

Spin-correlated radical pairs are transient reaction intermediates, often generated by photoinitiated, rapid electron transfer from a donor (in biological systems, often an aromatic amino acid) to an acceptor (here the FMN). As electron transfer occurs under conservation of total spin angular momentum, the total spin of the radical pair is defined by that of its molecular precursor (either a singlet |S⟩ (S = 0) or a triplet state |T⟩ (S = 1), where S is the total spin quantum number). If the radicals in the pair are weakly coupled the spin system is created in a superposition of the uncoupled states, consequently coherent interconversion occurs between the |S⟩ and |T⟩ states, driven by the interactions between electron and nuclear spins (such as ^1^H and ^14^N), [Fig. 1 B]. At zero and low static magnetic fields, this interconversion occurs rapidly involving all four states, but at higher fields, singlet-triplet mixing is restricted to |*S*⟩ and |*T*_0_⟩ with |*T_±_*⟩ energetically isolated from |*S*⟩ and |*T*_0_⟩ by Zeeman splitting [24, 25]. Importantly, singlet and triplet radical pairs have different fates; whilst the singlet pair is able to recombine to yield the ground state donor and acceptor, the triplet cannot do so but instead forms other forward photoproducts (e.g., by protonation/deprotonation), a route also and additionally open to the singlet pair. Under continuous illumination, the impact of the field on the singlet-triplet interconversion can be detected conveniently if either donor or acceptor or both form fluorescent excited states. In this case, a drop in fluorescence with applied magnetic field is expected for a triplet-born pair (impeded to mix into the recombining singlet state). Conversely, a resonant field reconnects the |*S*⟩/|*T*_0_⟩ manifold with |*T_±_*⟩, leading to an increase in fluorescence intensity. Therefore, we undertook MFE and ODMR measurements to test these hypotheses and hence to confirm the radical pair mechanism of MagLOV is triplet-born [24].

Both MFE and ODMR imaging were performed using a wide-field epifluorescence microscope (see S1.6, S1.7). Cells were grown in liquid culture, then washed and diluted into (non-autofluorescent) PBS buffer allowing us to image fields of single cells (S1.14). Samples were confined between two glass coverslips atop a stripline antenna printed circuit board (PCB) (see S1.13). The antenna PCB was placed inverted on the microscope stage and either an electromagnet (for MFE) or a permanent magnet (for ODMR) was positioned above. For MFE experiments, the electromagnet supplied a static field of 0 mT or 10 mT at the sample. For ODMR experiments, the permanent magnet supplied a static field (*B*_0_ ~ 20 mT) in the *z*-direction (S1.8), perpendicular to the radio-frequency (RF) field (*B*_1_ ~ 0.2 mT) supplied by the stripline antenna (S1.10). First, we measured the MagLOV MFE (see an example trace in Fig. 1 C), confirming that MagLOV exhibits a large MFE of Δℐ/ℐ_off_ = −50% where ℐ is the fluorescence intensity, ℐ_off_ is the fluorescence intensity immediately prior to switching the magnet on, and Δℐ = ℐ − ℐ_off_. In Fig. 1 D, we show the ODMR resonance of a single cell and the ODMR resonance averaged over many (~ 1000) single cells. For ODMR measurements, the detrended signal is Δℐ/ℐ_bg_, where ℐ_bg_ is a background curve fit to the data as described in S1.15.2, and Δℐ = ℐ − ℐ_bg_. As expected, the signal-to-noise is significantly improved by averaging over many cells in a field of view; however, it is also possible to extract an ODMR signal from a single cell, with an ODMR contrast of 10% [Fig. 1 D]. The remarkable per-cell magnetic sensitivity of *η*_0_ = 26 *µ*T Hz^−1/2^ (see S1.12) is afforded by a combination of optical detection and the high spin polarisation of the radical pair system [3]. Next, we recorded ODMR spectra at various static (*B*_0_) fields by adjusting the *z*-position of the static magnet [Fig. 1 E]. The central ODMR resonance follows the expected ESR relationship 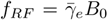, confirming the RF field is driving spin transitions of a 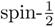 electron [52, 53]. Finally, we performed additional control experiments confirming the ODMR signal’s source, verifying against negative controls that an ODMR signal is present only when the MagLOV protein is expressed in S1.9.

## 3 Magnetic Field Effect can be Engineered by Directed Evolution

The AsLOV2 domain has been widely used as a starting point for engineering optogenetic and other light-dependent protein functionalities [54, 55]. For MFPs, depending on a target application (such as lock-in signal detection, multiplexing, or sensing) one could choose to optimise for metrics including MFE size, saturation rate (speed at which maximum MFE size is reached), ODMR contrast, ODMR saturation rate, and others; here we demonstrate this potential by performing selection to improve MFE contrast and saturation rate, generating variants specialised for applications demonstrated later in our work.

Starting from ancestor variant AsLOV2 C_450_A [9, 56] we used directed evolution to create variants of MFPs (summarised in S1.24). This engineering process (S1.3) involved successive rounds of mutagenesis (introducing all single amino acid changes to a given variant), followed by screening of samples from this variant library to select for increased MFE magnitude, eventually yielding MagLOV 2. To demonstrate the possibility of selecting on another metric using the same methodology, we further engineered MagLOV 2 by selecting for maximisation of saturation rate (rather than magnitude) of MFE, producing MagLOV 2 fast. We chose four variants to characterise in detail, which was done both using our microscope setup as described above [Fig. 2 A], and measurement of bulk cell suspensions [Fig. 3 and SI Fig. S11 A].

**Fig. 2.**
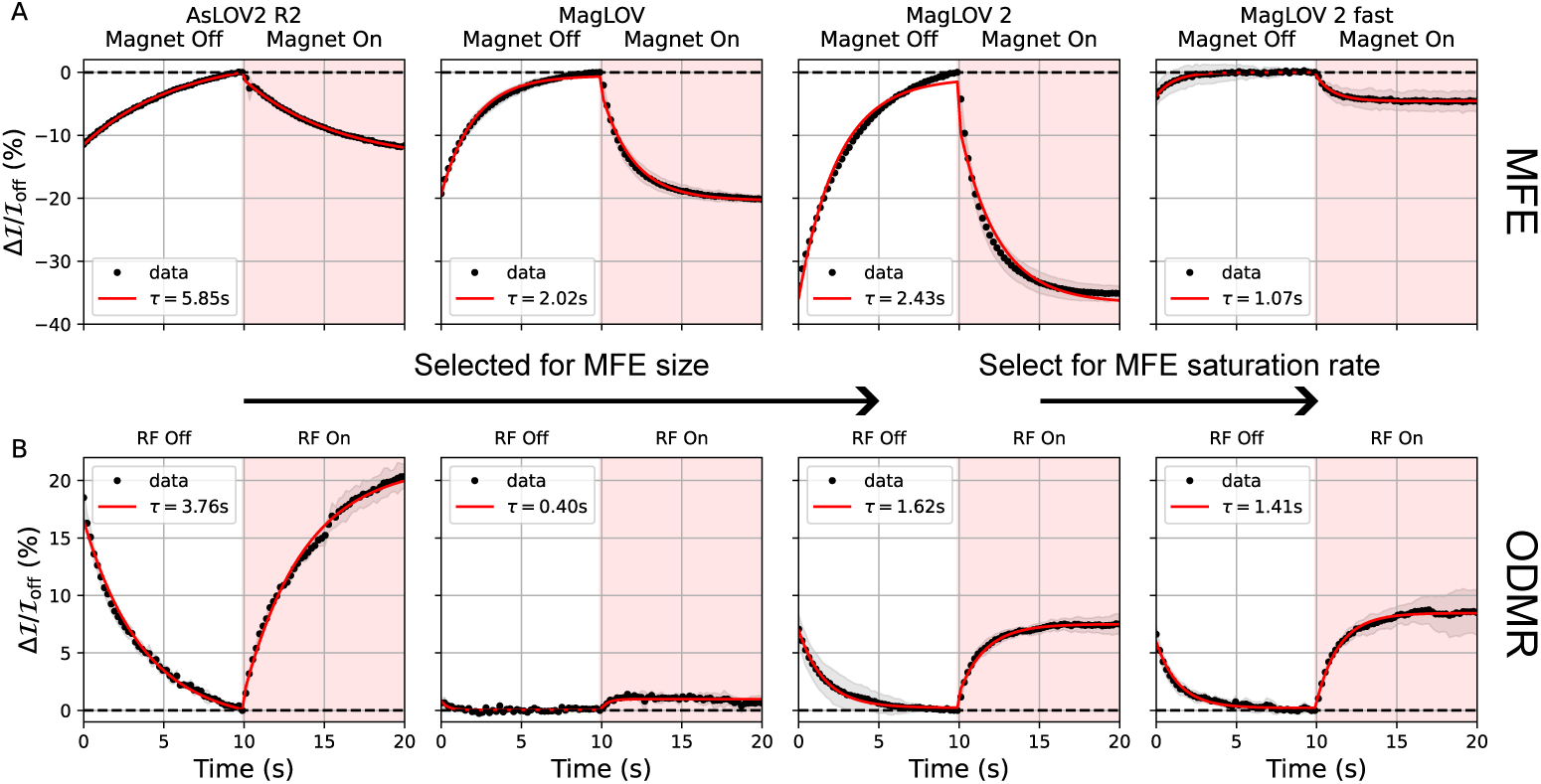
MFE and ODMR Dynamics of Engineered Variants. **A.** MFP variants were engineered by mutagenesis and directed evolution; from AsLOV2 R2 to MagLOV2 selection was performed to increase the MFE at saturation, while MagLOV 2 fast was selected for increased rate (i.e. reduced *τ* when the MFE step is modelled as *e^−t/τ^* [39], as shown in red). In these measurements a magnetic field of *B*_0_ = 10 mT was switched on and off with a period of 20 s. The traces shown are the average (and standard deviation, shaded) over multiple periods - as a result, some of the error in the standard deviation results from the background curve fit which attempts to match the true background curve (see S1.15.2 for details). **B.** A similar experiment was performed using ODMR on resonance, with a constantly applied static field of *B*_0_ = 21.6 mT and corresponding resonant field *B*_1_ frequency of *ω_RF_* = 604 MHz switched on and off with a 20 s period, averaged as above. Both sets of experiments were performed in identical imaging setups (with the sample on the RF antenna for A. to ensure the same lighting conditions as B.) with an illumination intensity of ~ 800 mW/cm^2^ at 450 nm. Note that *B*_0_ in A. is half that of *B*_0_ in B., explaining why the ODMR contrast is in some instances larger than the MFE.

**Fig. 3.**
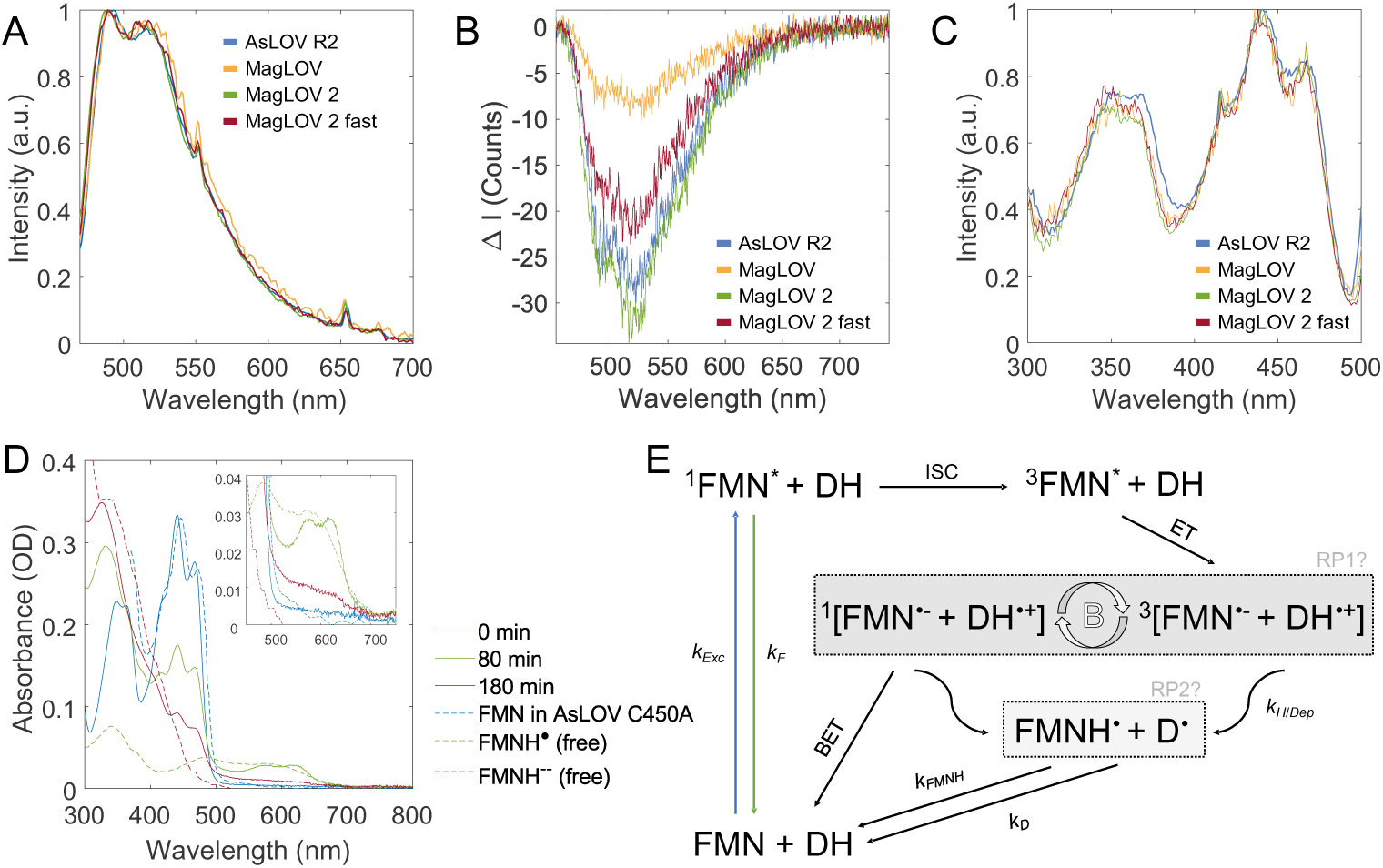
Spectroscopic characterisation evidences bound flavin-based radical pair. **A.** Normalised emission spectra with 450 nm excitation acquired for cell suspensions in PBS buffer. The spectra have been smoothed with a moving average filter over a ~ 1 nm bandwidth. **B.** Wavelength dependence of magnetic field-induced change of emission intensity where Δℐ = ℐ(*B*_0_ = 10 mT)−ℐ(*B*_0_ = 0 mT). Note that here we display the absolute intensity change as division by ℐ(*B*_0_ = 0 mT) obscures the wavelength dependence. A moving average filter over a ~ 1 nm bandwidth was applied to the spectra. **C.** Normalised excitation spectra for 510 nm emission also acquired for bulk MagLOV cell suspensions in PBS buffer. Vibrational fine structure in the *S*_0_ → *S*_1_ and *S*_0_ → *S*_2_ bands (where *S_n_* is the *n*^th^ excited singlet state) centred at ~ 450 nm and ~ 350 nm, respectively, indicates the emitting flavin is bound. **D.** UV-Vis absorption spectrum of purified MagLOV 2 fast at different times after the onset of blue LED illumination. Inset shows a zoomed-in wavelength region with the characteristic absorption of FMNH*^•^* (green). Literature reference spectra are shown in dashed lines [57–59]. **E.** Proposed photoscheme. Following photoexcitation (*k*_Exc_), the excited singlet flavin (^1^FMN*) can either emit a photon (*k*_F_) or undergo intersystem crossing (ISC) to the excited triplet state (^3^FMN*). The primary radical pair (RP1) is formed by electron transfer (ET) from a nearby donor, and can undergo singlet-triplet interconversion, which is altered in the presence of an applied magnetic field (*B*). Only the overall singlet RP can undergo back-electron transfer (BET) to reform the ground state, while either RP can form secondary (perhaps spin uncorrelated) radicals (RP2) through protonation and/or deprotonation reactions (*k*H/Dep). These long-lived secondary radicals return to the ground through slow redox reactions (*k*FMNH, *k*D).

The observed differences in MFE can be interpreted based on past work investigating flavin magnetic field-sensitive photochemistry: MFE enhancement kinetics (i.e. time to MFE saturation) are determined by the ratio between the rates that donor and acceptor free radicals return to the ground state [39]. With the mutations introduced in MagLOV 2 fast, the time constant of the response is decreased, which may indicate the acceptor return rate increases relatively to the donor. Interestingly, we observe the ODMR contrast and rates of each variant differ significantly [Fig. 2 B], but not necessarily in simple correlation with MFE magnitude. This raises the possibility of engineering orthogonal fluorescent signatures, expanding again the number of tags available for multiplexing - for instance, with further engineering the total might be (number of emission colours)×(number of resolvable MFE signatures)×(number of resolvable ODMR signatures).

## 4 Bound Flavin Radical Pair Model supported by MFE Spectroscopy

Adopting the SCRP model prompts consideration of the electron donor and acceptor identities. Both previous studies [46, 48], and our spectral data in [Fig. 3], support identifying the acceptor molecule as the FMN cofactor. For all variants of MFPs expressed in cells, we find that the wavelength-resolved fluorescence intensity modulated by the applied magnetic field (Δ*I*) in [Fig 3 A, B] matches the FMN emission spectrum [56, 60], supporting that both MFE and ODMR are detected on the flavin emission. The excitation spectrum shows vibrational fine structure [Fig 3 C] and is in excellent agreement with the dark-state absorption spectrum of AsLOV2 C_450_A [57], confirming the emission originates from *bound* FMN. Furthermore, this corroborates control experiments (SI S1.9) that show observed ODMR signatures are not a result of cellular autofluorescence. The absorption spectrum of purified MagLOV 2 fast, before and after continuous irradiation with blue light, is shown in Fig. 3 D (full temporal evolution is provided in Fig. S11 D). The slow formation of the stable radical FMNH*^•^* after several minutes of illumination is characterised by the appearance of a broad band featuring two peaks centred around 575 nm and 615 nm, as observed for AsLOV C_450_A, and accompanied by the expected decrease in ground state absorption, centred around 450 nm [57]. At later times, most of the flavin is converted to the fully reduced form, FMNH^−^, resulting in a rise in absorption around 325 nm, while some remains present as FMN and FMNH*^•^*. The blue shift in MagLOV-bound FMN absorption relative to AsLOV C_450_A, evident both in the excitation and absorption spectra, may indicate a change in polarity of the flavin binding pocket. The observed photostability of MagLOV (Figure S11 B) correlates with the slow formation of FMNH*^•^* on a minutes timescale in these conditions. In comparison, related flavoproteins, such as cryptochrome, are reduced to the semiquinone state within seconds even under much weaker irradiation intensities [61]. Unlike the recently reported MFE in mScarlet3 and FMN mixtures, which rely on a bimolecular reaction between the excited state mScarlet3 and a fully reduced flavin in solution [41], MagLOV requires no pre-illumination or additives for an MFE to develop as the FMN is non-covalently bound.

The formation of FMNH*^•^* from the excited triplet state, ^3^FMN*, likely proceeds via initial formation of FMN^•−^ in a SCRP on a nanosecond timescale as in AsLOV C_450_A [46], followed by slow protonation. Alternatively, FMNH*^•^* could be formed by proton-coupled electron transfer. A proposed photoscheme is given in Figure 3 E, and a model based on this photoscheme is simulated in SI S1.18, which successfully fits both the experimental MFE and ODMR data. In cryptochrome, the SCRP is formed by a cascade of ETs along a tryptophan tetrad [43], however, a single donor aromatic amino acid can be sufficient, as demonstrated by the magnetosensitivity of cryptochrome-mimicking flavomaquettes [62, 63] and FMN bound inside the bovine serum albumin (BSA) protein[64]. Regarding the donor species, single point mutations in AsLOV2 C_450_A leading to quenching of the emissive NMR signal suggest W_491_ as the electron donor [65], which was corroborated by isotopic labelling of Trp residues [66]. However, given the extent of mutations in the variants studied here, we cannot confirm W_491_ is still the counter-radical. In the structurally related iLOV-Q_489_, derived from *A. thaliana* phototropin-2 (AtPhot2) LOV2 domain, transient absorption spectra revealed a neutral tryptophan radical, Trp*^•^*, is formed in conjunction with FMNH*^•^*, and photoinduced flavin reduction in single-point mutations of selected tyrosine and tryptophan residues suggested several amino acids might be involved in SCRP formation [67].

## 5 Applications of Magneto-Sensitive Fluorescent Proteins

### 5.1 Demonstration of Multiplexing and Signal Lock-in

Our library of MFP variants exhibits differences in the rate and magnitude of response across both MFE and ODMR characterisation [Fig. 2]. Where such differences can be engineered orthogonally between variants, they open the possibility of using libraries of MFP reporters as a multiplexing tool to extract several signals from a single measurement modality. To demonstrate this potential in fluorescence microscopy, we first cloned strains of AsLOV R5 (one of the variants evolved between AsLOV R2 and MagLOV, see S1.24) and MagLOV 2 into high constitutive expression strains (SI S1.1). We initially characterised each variant in isolation, measuring fluorescence traces for ~ 2000 cells in a field-of-view [Fig. 4 A]. The same processing was performed as in Fig. 2 but to individual cells, yielding a value of *τ* (the MFE saturation timescale) for each cell, which allows measurement of each sample’s intrapopulation variability [Fig. 4 B]. We then characterised a mixture of these variants (again ~ 2000 cells) [Fig. 4 C], for which parameter values fit to single cells show strong bi-modality, enabling the population to be decomposed into sub-populations identified by which magneto-sensitive fluorescent variant is expressed. To perform the decomposition, we trained the machine learning classifier XGBoost [68] on the dynamic data (i.e. fluorescence versus time) used to generate Fig. 4 B, and (without further training) this was used to classify the cells in Fig. 4 B. The ratio of the two variants by classification was R5 : ML2 = 0.9 : 1, which likely differs from the anticipated 1 : 1 ratio both due to classifier accuracy as well as typical measurement errors expected from use of OD600 to quantify cell counts [69]. Unlike the data shown in Fig. 2, here all single cell MFE traces were normalised (on a per-cell basis), meaning the classifier only utilises the relative shape of the curves and not the magnitude of the MFE, and so is robust to noise or offset in the absolute brightness of the signal, as might be caused by scattering or autofluorescence (which pose practical limits on many sensing applications as described in S1.22). Future application of this sub-population labelling technique would benefit from engineering of variants with greater variation in dynamics such that the separation of histograms (as in Fig. 4 B,C) is significantly greater than each population’s intracellular variability.

**Fig. 4.**
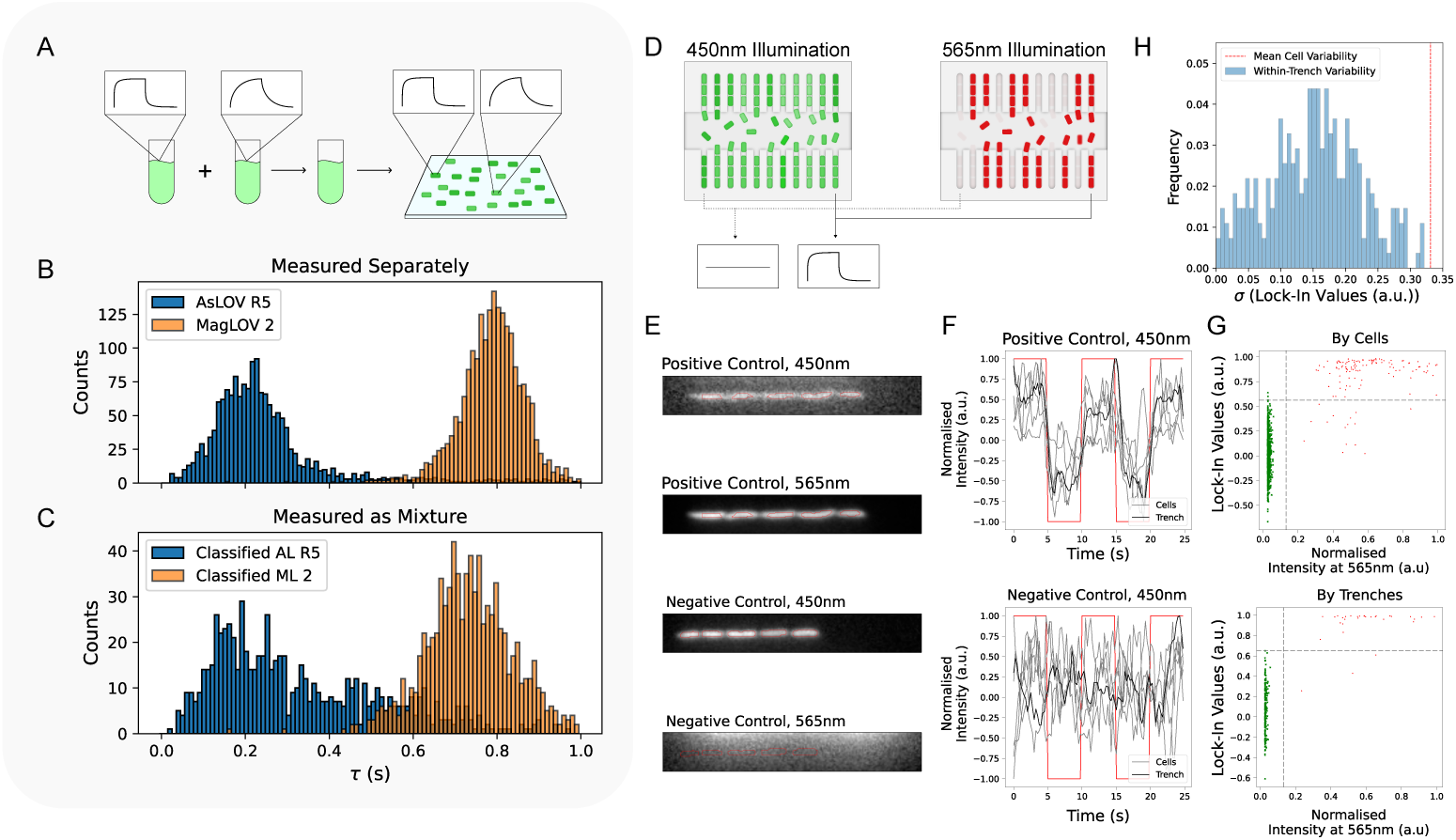
Multiplexing and Lock-in Applications using MFE. **A.** Illustration of the preparation of multiple variants on a coverslip. For this application, AsLOV R5 and MagLOV 2 were expressed in MG1655 cells using a strong RBS and constitutive promoter (see SI S1.1). **B.** Exponential curves with timescale parameter *τ* were fit to the MFE of each cell in each field-of-view. In B measurement of separate populations illustrates MagLOV 2 has a greater MFE saturation timescale *τ* than that of an earlier variant AsLOV R5. **C.** Next, the populations were mixed in an equal ratio (determined by OD600). A classifier (XGBoost [68]) was trained on the time-series data used to create B, and then used to classify the data in C. Prior to training, and for classification, the MFE response (Δℐ/ℐ_off_) curves (sketched in A) of each cell were normalised to range from 0 to 1. **D.** Schematic illustration of the microfluidic setup. For the purpose of demonstrating the utility of lock-in in this experiment, cells were engineered to weakly express MagLOV and to co-express mCherry as a control. These cells were mixed with cells expressing only EGFP, such that we could demonstrate isolating MagLOV cells from other, non-magnetic responsive fluorescent reporters in the same spectral range, and under conditions where MagLOV produces only a small signal. The microfluidic chip is composed of two rows of trenches (vertical lines) fed by a central channel bringing fresh media into the system and carrying excess cells out. **E.** Cropped view of single trenches, with cells circled in red as identified by a cell-segmentation algorithm (S1.16). **F.** Time-series of the 450 nm illumination fluorescence response for the individual cells depicted in E over time, as a magnetic field *B*_0_ = 10 mT is switched on and off (red line). Both the traces for each cell in the trench (grey lines), and the average trace over all the cells in the trench (black line) are shown. **G.** Confusion scatter plots for classifying whether cells express MagLOV or EGFP based on magnetic response, taking the mCherry 565 nm fluorescence (red) as ground-truth and using the MFE lock-in value (S1.16) (green) as the predictor. Top left quadrant are false negatives, top right quadrant are true positives, bottom left quadrant are true negatives, and bottom right quadrant are false positives. The balanced accuracy by cells and by trenches is 0.99, see S1.23 for classification data. **H.** The standard deviation *σ* of the lock-in values in G is calculated between cells in each trench (blue histogram) and between all cells in all trenches (mean value in red dashed line).

Using a microfluidic setup (see S1.5), we further investigated the intracellular variability of MFP variants, and demonstrate the possibility of lock-in detection in weak signal environments. Cells expressing EGFP [70], and cells co-expressing mCherry [71] with very weak MagLOV expression (predicted MagLOV expression levels ~ 0.04% of that in multiplexing experiments above, SI S1.4 for details) were mixed and loaded into a microfluidic “mother machine” chip consisting of evenly spaced trenches, which after a few hours of growth are each filled only with cells whose ancestor is the cell at the closed end of the trench (i.e. they may be considered clonal) [Fig. 4 D]. The cells were imaged using the same widefield fluorescence microscopy setup as described previously. Using MFE-based lock-in detection [64], we found that MagLOV cells could be identified distinctly from EGFP with balanced accuracy ~ 0.99 (using mCherry as a ground truth), with accuracy improving when averaging over a trench compared to distinguishing single cells [Fig. 4 E-G]. This approach allows variability to be attributed to inter-clonal or intra-clonal sources; we observe that mean intra-trench variability (quantified by standard deviation over 5-8 cells in the trench) is approximately half that of the variability over all MagLOV-positive cells [Fig. 4 H]. This suggests approximately one quarter of the total variance arises from intra-clonal noise sources (e.g. phenotypic variability over 2-3 generations, camera and accompanying measurement noise), with inter-clonal sources (longer term phenotypic variability, variation in local environment) contributing the remaining three quarters.

### 5.2 Spatial Localisation

Methods for the spatial localisation of fluorescent signals in biological samples such as cell cultures and tissue samples are of significant interest for both diagnostics and treatment development [72, 73]. However, techniques based on localisation using fluorescence, for example fluorescent modulated tomography (FMT), are challenging due to the inherent scattering and absorbing nature of tissue, and the requirement to localise the fluorescence via detailed modelling and inversion of the optical signal [74, 75]. As such, we sought to explore whether MagLOV could be employed as a fluorescent marker localised by optically detected magnetic resonance imaging.

First, we tested localising MagLOV in the widefield microscope setup, using a permanent magnet to vary the resonance condition across the field of view [Fig. 5 A]. The sequence of images acquired during a RF *B*_1_ frequency sweep was subsequently integrated over cross-sections in which the *B*_0_ field is approximately constant, yielding an image [Fig. 5 B] where the frequency of peak response (y-axis) denotes the position in space along the *z*-axis, which varies across a 0.5 mm field-of-view as anticipated for a field gradient of 1 mT/mm.

**Fig. 5.**
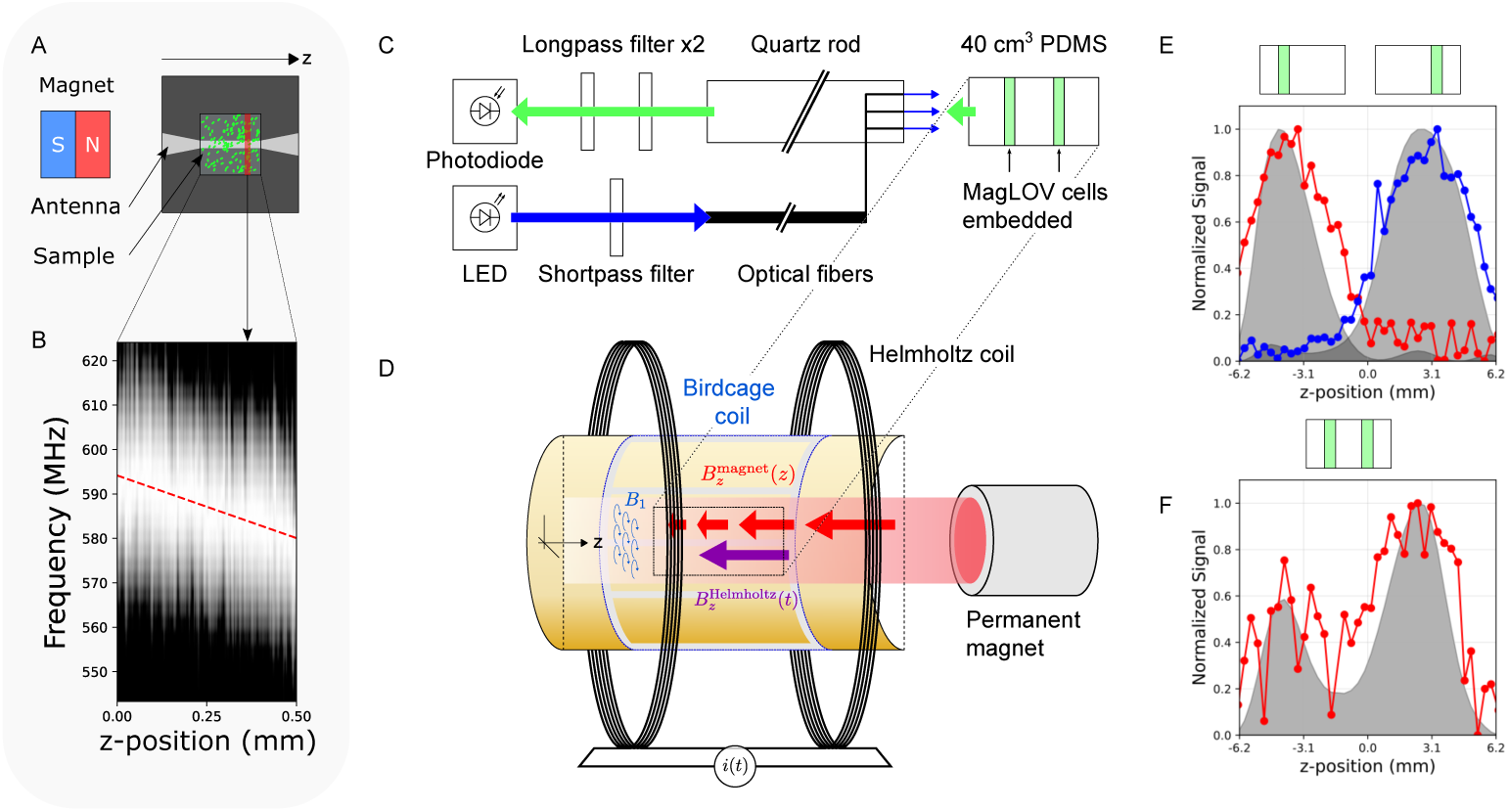
Spatial Localisation using ODMR. **A.** Schematic of the widefield setup for demonstrating localisation of cells in a 2D plane. The permanent magnet creates an approximately linear gradient field *B*_0_ oriented in the *z*-axis, that is perpendicular to the radio-frequency *B*_1_ field (which rotates at the Larmor frequency radially around *z*) generated by the RF coil. The radiofrequency of *B*_1_ is scanned while the entire field-of-view is imaged. **B.** Subsequently, the images are divided into regions (e.g. highlighted in red in A.) and integrated over that region. The integrated brightness vs. frame number (i.e. frequency) forms a 1-dimensional spatial map - in this case the sample is present in the entire field of view thus we see a diagonal line, shifting by ~ 14 MHz over the 0.5 mm field of view as anticipated for the 1 mT/mm field gradient. **C, D.** Schematic of the custom-built MRI setup illustrating the optical illumination and collection paths (C.), and the magnetic field gradient inside the resonant MRI coil (D.). See Fig. S17 for photographs. **E, F.** We embedded cells expressing MagLOV 2 fast (chosen for its ODMR contrast, fast saturation, and low overall brightness to make detection challenging) into a PDMS sample at two different positions along the coil axis, separately (E.) and simultaneously (F.). Lock-in ODMR detection was used to locate the samples along the coil axis. The red and blue curves are the measured data, and grey shaded regions are after processing that data with a de-convolution. The location of the samples is clearly identifiable on their own, and resolvable via deconvolution together. Against a ground truth separation of 7.5 mm, and assuming the measured field gradient of 0.95 mT/mm is uniform within the coil, deconvolved individual samples had a calculated peak separation of 6.6 mm and the combined sample 6.1 mm. Using the individual sample data to calibrate this measurement yields a two-sample distance estimate of ~6.9 mm. Note that the de-convolution makes no assumptions about the number or location of peaks, only an estimated point spread function is required as input – which is the known ESR linewidth here.

Next, we converted a preclinical MRI 28 mm diameter “Birdcage” RF coil, used for creating a spatially highly homogeneous *B*_1_ RF field at 500 MHz, into a novel optically detected fluorescence MRI instrument via integration of a fibre-coupled illumination system and imaging using a photodiode ([Fig. 5 C,D] and S1.20). Note that the photodiode is in effect a ‘single-pixel’ detector, meaning it collects no spatial information and directional scattering of light would cause no reduction in information of final signal (apart from a possible decrease in absolute brightness). We used a permanent neodymium rare-earth magnet to impose a static magnetic field, *B*_0_, which varied linearly along the *z*-axis with a gradient strength of approximately 0.95 T m^−1^ at the RF isocentre of the coil used. The imaging isocentre is both spatially the centre of the RF coil’s sensitive area and was chosen to be at 17.8 mT corresponding to an ESR Larmor frequency of *f* ~ 500 MHz, the coil’s resonant frequency. The magnetic field was modulated axially by a Helmholtz coil such that 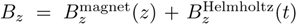 where *t* is time. By varying the Helmholtz coil current between ±1 A, we were able to shift the effective magnetic field by ±5.87 mT around the resonant condition, providing a total field-of-view of approximately 12 mm along the gradient direction (see S1.20 for a detailed calculation). The device is therefore able to scan spatially in one dimension as the Larmor frequency remains fixed at 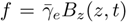, where different spatial positions are brought into resonance through current modulation while maintaining fixed RF excitation.

As the *B*_0_ field is swept, the *B*_1_ field is switched on and off such that the ODMR contrast can be detected via lock-in detection. To simulate a 3D volumetric sample, we cast a 40 cm^3^ volume of PDMS (matching the region of uniformity of the RF coil) with empty cylinders of 0.4 mm diameter embedded in it perpendicular to the *B*_0_ gradient and separated in *z* by 7.5 mm. We filled these cylinders with cells expressing MagLOV 2 fast, measuring both at different depths individually and together; we found that good localisation could be achieved when isolating a single sample (Fig. 5 E), and despite an increase in noise, deconvolution of the signal enabled two cylinders at different depths to be simultaneously localised (Fig. 5 F) to within ~ 0.6 mm of a calibrated ground truth. We note that the generation of homogeneous RF fields at ~ 500 MHz within living systems (including humans) is an active area of research with known working designs for different species and anatomies [76–78], and that the continuous-wave nature of this experiment avoids the need to undertake a pulsed ESR experiment within the short relaxation lifetimes inherent to spin-correlated radical pairs at high temperature. In this way, our approach forms a novel alternative method for spatially-resolved and scattering-insensitive sensing of genetically-encodable fluorescent proteins.

### 5.3 Microenvironment Sensing

We next sought to determine whether MagLOV could function as a quantum sensor of its local environment. Although FMN is bound within the protein scaffold, we hypothesised that it might exhibit magneto-optical properties that depend acutely on nearby paramagnetic species, analogous to nitrogen-vacancy (NV) centres in nanodiamond-based sensing approaches [4, 79, 80]. To test this we diluted purified MagLOV 2 fast samples to 3.7 *µ*M in solutions containing the paramagnetic contrast agent Gadolinium (total spin *S* = 7/2; Gd^3+^ chelated as the MRI contrast agent Gadobutrol). We anticipated that these freely diffusing paramagnetic species would act like diffusing point dipoles, modulate the dipolar interactions of the radical pair, and lead to enhanced relaxation and characteristic attenuation of the MFE contrast. This hypothesis was confirmed by our measurements which show that, despite the constrained cofactor geometry, MagLOV exhibits a clear dose-dependent reduction in MFE signal with increasing paramagnetic ion impurity concentration. We note that there was no visual change in the microscopy images, nor any Gadobutrol concentration-dependent difference in absolute brightness of the sample, indicating the chelated Gadolinium did not cause protein denaturation or aggregation, or FMN dissociation. More quantitatively, paramagnetic impurities can be modelled as stochastic point dipoles that modify the *T*_1_ or *T*_2_ relaxation rates in radical pair kinetics [81], thereby changing the contrast. We therefore anticipated that a characteristic timescale for this process, *τ*, would scale as 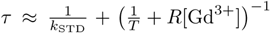 where *k*_STD_ is a stochastic decoherence rate, *T* is a semiempirical *T*_1_ or *T*_2_ relaxation time, and *R* the effective relaxivity of the paramagnetic impurity. For a freely diffusing paramagnetic impurity, we expect the normalised spectral density function to contain a molecular rotational contribution term ~ *k_B_T/*8*πd*^3^*η* from Stoke’s law, and therefore, we expect contrast to fit a functional form ~ (*a*[*x*] + *b*)^−1^+*c*, and indeed this is matched in Fig. 6. Overall these results imply MagLOV can be used as an *in situ, in vivo* quantum sensor which, analogous to other quantum sensing modalities, opens possible applications as a sensor of the cellular microenvironment.

**Fig. 6.**
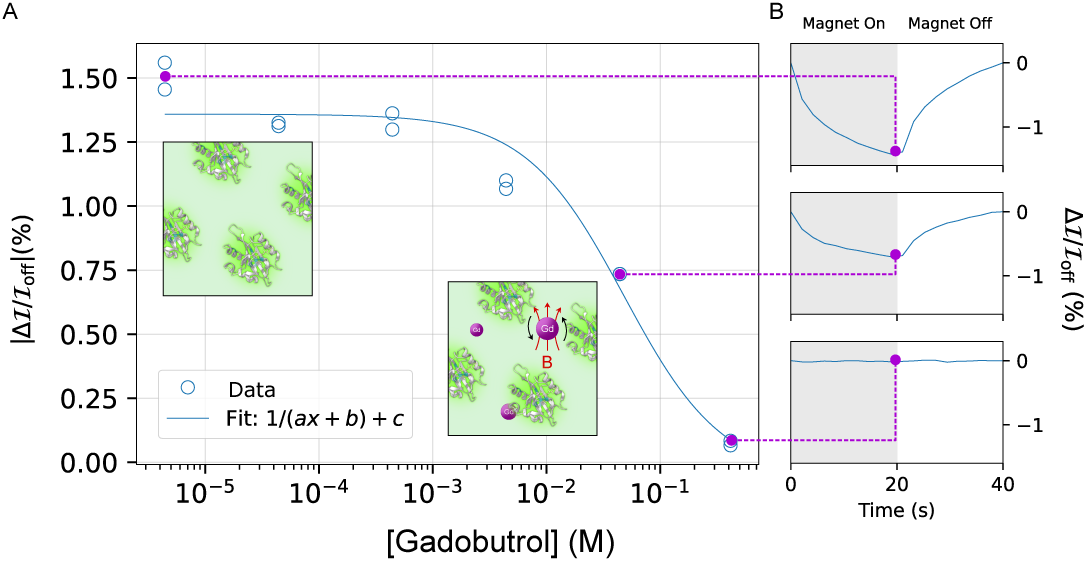
Local Spin Environment Sensing using MFE. **A.** The MFE contrast of purified MagLOV 2 fast is measured at varying concentrations of the paramagnetic contrast agent Gadobutrol (see S1.21 for experimental setup), demonstrating an reciprocal dependence on concentration, consistent with spin relaxation and indicating that MagLOV is sensitive to its surrounding spin environment. **B.** The MFE time trace is shown for three points on the dose dependence curve of A. In general, we observed significantly lowered MFE for purified protein than in cells, see SI S1.12 for further data.

## 6 Discussion

Directed evolution, enabled by straightforward fluorescence screening, has proved to be a powerful technique to engineer proteins exhibiting magneto-sensitive responses. The advent of stable, highly responsive magneto-sensitive proteins represents a paradigm shift; transitioning from quantum biological systems studied primarily for scientific interest toward engineerable tools with potential for widespread application. Previously existing natural and engineered proteins (typically designed as model representatives of the cryptochrome) exhibited comparatively small responses to magnetic fields, typically required sophisticated experimental apparatus for study, did not exhibit measurable MFEs in living cells, were prone to rapid light-induced degradation, and were therefore unsuitable for biotechnological applications or high-throughput setups required for directed evolution [37, 40, 62]. Furthermore, compared with other candidates for quantum biological sensing, two unique advantages of a protein-based system are (a) that it is configurable: significant engineering improvements can be made (relatively simply) by changing the DNA encoding the protein, and (b) that it can be endogenously expressed, enabling coupling of their expression to diverse genetic or chemical signals. MFPs therefore are the best of both worlds, enabling sensitive quantum measurements while also being highly amenable to engineering and cellular integration.

Using this system for magnetic resonance measurements enables a host of applications. First, modulating MagLOV fluorescence by applying a time-varying magnetic field enables multiplexing and lock-in detection. Multiplexing can enable dramatic expansion in the number of fluorescent reporters that can be distinguished in a single experiment. Meanwhile, lock-in could enable fluorescent protein measurements to be performed where previously not possible due to poor signal quality, for example if only small quantities of a fluorescent marker can be produced, or in tissue measurements where autofluorescence and scattering are limiting factors [26, 64, 82, 83]. Second, we demonstrate that using ODMR it is possible to spatially localise fluorescence signals in a three-dimensional volume, utilising the fact that resonance only occurs when the required conditions are met by (orthogonally controllable and tissue-penetrating) RF and magnetic fields. Finally, magnetic resonance sensing can be used to determine the presence of molecular species creating local magnetic noise in our protein’s environment. This could be used to measure the presence of free radicals or paramagnetic metalloproteins, both critical to a number of physiological processes [80].

While the properties of the MFPs we generated are far superior to previously studied proteins that exhibit MFEs (S1.17.4), their optimisation is by no means complete. Much like fluorescent proteins, we expect MFPs may be engineered to make general improvements, such as to solubility, photostability, spectral response, and quantum-yield (see S1.17.4), as well as further improving their MFE/ODMR properties. There also remains significant opportunity for mechanistic investigation utilising the wide array of techniques previously applied to biological systems exhibiting MFEs and ESR [40, 44, 47, 55, 84, 85]. Crucially, mechanistic understanding and high throughput bio-engineering (similar to and more advanced than we demonstrate here) can go hand in hand - for instance, enabling the creation of rational design tools that can optimise MFE/ODMR properties for a specific application. Finally, we hope that the development of MFPs can serve as the starting point for a novel class of magnetically controlled biological actuators, whereby application of a local magnetic field can be be coupled to downstream cellular effects – such a technology would be of significant biomedical and biotechnological interest.

## Supplementary information

- SI.pdf

## Acknowledgements

The authors would like to thank Kevin Henbest, Emil Vatai and Luca Gerhards for helpful discussions, Daniel Cubbin for help with the UV-Vis characterisation, and Islay Robertson and Phillip Reineck for assistance with proof of concept experiments. Carolyn Carr and Greg Mazur graciously lent us RF equipment. The protein structure in Fig. 1 was rendered using ChimeraX [86].

The EcoFlex kit was a gift from Paul Freemont (Addgene kit #1000000080) [87].

## Declarations

GA and SS were supported by funding from the Biotechnology and Biological Sciences Research Council (UKRI-BBSRC) [grant number BB/T008784/1]. JJ and VTF were supported by funding from the Engineering and Physical Sciences Research Council (UKRI-EPSRC) [grant number EP/W524311/1]. IK and HS are supported in part by the UKRI-EPSRC under the EEBio Programme Grant, EP/Y014073/1, and EP/X017982/1. RH is supported by funding from the UKRI-EPSRC [grant number EP/Y034791/1] AŠ, LMA and CRT are supported by the European Research Council under the European Union’s Horizon 2020 Research and innovation programme, Grant Agreement No. 810002, Synergy Grant: ‘QuantumBirds’, CRT thanks the US Army. JJM would like to acknowledge support from the Novo Fonden (NNF21OC0068683). HS recognises support from the Philip Leverhulme Prize.

## Data availability

The experimental data and that support the findings of this study are available in Zenodo with the identifier 10.5281/zenodo.14500235. Plasmid sequencing data have been made available on the European Nucleotide Archive (ENA) with the identifier PRJEB83586. Protein sequences have been submitted to UniProt and will be made available pending review.

## Unique biological materials

All biological materials (bacterial cells, plasmids) are available upon request.

## Author contribution

- Gabriel Abrahams - conceptualisation, development of ODMR setup, development of widefield MFE setup, experimental measurements, analysis, interpretation
- Vincent Spreng - sample preparation, widefield MFE measurements, cloning plasmids and strains (post directed evolution) strategy and execution
- Ana Štuhec - photochemical and spectroscopic characterisation, bulk MFE measurements, analysis, interpretation
- Robin Henry - sample preparation, widefield MFE and ODMR measurements, cloning plasmids and strains(post directed evolution) strategy and execution, purified protein sample quantification
- Idris Kempf - microscope platform development, performing microfluidics experiments, analysis for microfluidics experiments
- Jessica James - microscope platform development, carried out performing microfluidics experiments, analysis for microfluidics experiments
- Kirill Sechkar - microfluidics platform development, performing microfluidics experiments
- Scott Stacey - cloning plasmids and strains (post directed evolution) strategy and execution, figure preparation
- Vicente Trelles-Fernandez - cloning plasmids and strains (post directed evolution) strategy and execution
- Lewis M. Antill - spin dynamics simulations, analysis, interpretation
- Christiane R. Timmel - project co-ordination, analysis, supervision
- Jack J. Miller - design and development of imaging experiments, MRI hardware, analysis, interpretation.
- Maria Ingaramo - variant generation by directed evolution
- Andrew York - conceptualisation of MFE experiment, project co-ordination
- Jean-Philippe Tetienne - preliminary experiments, analysis
- Harrison Steel - project co-ordination and conceptualisation, development of microscopy setup, supervision

