## Supplementary Information for "Quantum Spin Resonance in Engineered Magneto-Sensitive Fluorescent Proteins Enables Multi-Modal Sensing in Living Cells"

### S1 Supplementary Methods

#### S1.1 Strains

T7 Express *E. coli* (New England Biolabs - C2566) were used for MFE and ODMR experiments in Fig. 1, 2 and 3. *E. coli* MG1655 cells were used for multiplexing and mother machine lock-in experiments in Fig. 4, due to its effectiveness as a heterologous gene expression strain, and past optimisation of mother machine chip loading protocols for this host. MG1655 was also used for Fig. 5. Note that MagLOV variants characterised in this strain exhibit different rates of MFE response which we attribute to changes in expression levels and cellular environment.

#### S1.2 Media and Chemicals

TB auto-induction medium (Formedium - AIMTB0260) was used for growth of liquid cultures of T7 Express strains. LB medium (Formedium - LBX0102) was used for growth of MG1655 strains in liquid cultures and agar plates (Formedium - AGR10). M9 medium (Formedium - MMS0102, supplemented with 0.2g/L Pluronic F-127 (Sigma Aldrich - P2443-250G), 0.34g/L thiamine hydrochloride, 0.4% glucose, 0.2% casamino acids, 2mM MgSO4, 0.1mM CaCl2, 50 mg/L EDTA Na2.2H20, 5mg/L FeCl3, 0.84 mg/L ZnCl2, 0.13 mg/L CuCl2.2H2O, 0.05 mg/L CoCl2, 0.1mg/L H3BO3 and 0.016 mg/L MnCL2.4H2O) was used for mother machine experiments. Antibiotic stocks were prepared as follows: 100 mg/mL carbenicillin (Formedium - CAR0025) dissolved in 50% (v/v) EtOH and 20 mg/mL chloramphenicol (Fisher - 10368030) dissolved in 100% (v/v) EtOH; stocks were diluted 1000x for experiments.

#### S1.3 Directed Evolution of Magnetic Fluorescent Protein Variants

We previously reported the directed evolution of magneto-fluorescent proteins (MFPs) resulting in MagLOV in [1]. Directed evolution of MFP variants was initiated with AsLOV2 C450A [2], a protein originally isolated from the common oat *Avena sativa*. The mutagenesis protocol was based on methods developed previously [3]. In particular, we used 142 PCR reactions with NNK semi-random primer pairs (supplied by Eton Bioscience) to make all single amino-acid mutants of AsLOV2 (404-546) C450A at all locations, which produces a library containing 2982 protein variants, though (due to the pooled nature of the screen) it is likely that not all variants are present for each round. This library was transformed into *E. coli* strain BL21(DE3) and spread across $\sim$ 10 LB-agar plates for screening. After transformation, plates were left for two days at room temperature (25° C) as this was observed to lead to stronger magnetoresponses, which we hypothesise arises due to changes in the intracellular chemical environment. Screening used the same fluorescence photography system described in [3], with addition of an electromagnet (KK-P80/10 - Kaka Electric) below the sample, which was turned on/off every 15 seconds using an Arduino Uno while acquiring successive images. After each screen images are processed to identify the single colony (across all plates) with the largest fractional change in fluorescence between the magnet on/off conditions. The selected colony was picked (manually) from the corresponding plate, and used as the basis for the next round of mutagenesis and selection. Subsequent mutagenesis rounds used the *same* primers to avoid re-making the primer the library with each mutation (which we hypothesise selects against multiple consecutive nearby mutations), except before rounds leading to AsLOV R5, MagLOV R7, and MagLOV 2 at which point primers in the library that overlapped mutated sites were updated to the current variant's sequence. In total 11 rounds of mutagenesis were undertaken, all selecting for amplitude of magnetic field effect *with the exception* of round 11 (MagLOV 2 fast) where mutants from round 10 were selected based on fast response time (quantified as the mutant with the largest contrast measured in the first 100 ms following magnetic field change). After each round of mutagenesis, selected variants were sequenced using the Sanger method (supplied by Quintara Biosciences). The variants measured in this paper and some intermediaries are listed in Table S6.

#### S1.4 Plasmid construction

Whole Plasmid Sequencing was performed by Plasmidsaurus using Oxford Nanopore Technology with custom analysis and annotation.

Derivative plasmids of those generated by directed evolution (for example MagLOV+mCherry) were constructed using the EcoFlex kit following standard EcoFlex protocols [4]. Level 1 assemblies were performed using NEB BsaI-HFv2, NEB T4 DNA Ligase Reaction Buffer (B0202S) and Thermo Scientific T4 DNA Ligase (EL0011). Level 2 assemblies were performed using NEBridge Golden Gate Assembly Kit BsmBI-v2 and NEB T4 DNA Ligase Reaction Buffer (B0202S).

Level 1 PCR cycling protocol: 30x(5min@37° $\rightarrow$ 5min@16°) $\rightarrow$ 10min@60° $\rightarrow$ 20min@80°C

Level 2 overnight PCR cycling protocol: 65x(5min@42°C $\rightarrow$ 5min@16°C) $\rightarrow$ 10min@60°C $\rightarrow$ 20min@80°C

*pGA-01-01* An EcoFlex Level 1 part with T7 promoter was manufactured by Twist Bioscience. An EcoFlex Level 1 reaction was performed to integrate the part into the EcoFlex pTU1-A-RFP backbone. The plasmid was transformed into NEB Dh5$\alpha$ *E. coli* for plasmid production and into NEB T7 Express *E. coli* for experiments.

*pTU1* We used pTU1 as a negative control plasmid [Fig. S4] which comes direct from EcoFlex kit. The plasmid was transformed into NEB Dh5$\alpha$ *E. coli* for plasmid amplification and into NEB T7 Express *E. coli* for experiments.

*pRSET-AsLOV_R2, pRSET-MagLOV, pRSET-MagLOV-2, pRSET-MagLOV-2_R11-f* Plasmids containing magnetic fluorescent variants under control of T7 promoter were created by directed evolution as described in Section S1.3. *E. coli* NEB T7 were transformed for measurements. The predicted translation rates for these strains is $50 \times 10^3$[5]

*pVS-01-03_EGFP* This EGFP expressing plasmid was assembled solely from EcoFlex parts (pTU1-B-RFP, pBP-J23108, pBP-pET-RBS, pBP-ORF-eGFP, pBP-BBa-B0012). It was used for mother machine lock-in experiments.

*pVS-02-04_mCherry+MagLOV* EcoFlex Golden Gate assembly was used to create a plasmid that expresses both MagLOV and mCherry. In a level 1 reaction, pVS-01-01_mCherry and pVS-01-02_MagLOV were assembled using the following bioparts from the EcoFlex kit. pVS-01-01_mCherry: pTU1-A-lacZ, pBP-J23108, pBP-pET-RBS, pBP-ORF-mCherry, pBP-BBa_B0012. pVS-01-02_MagLOV: pTU1-B-RFP, pBP-J23119, pBP-pET-RBS, BP-01_MagLOV, pBP-BBa_B0012. The BP-01_MagLOV sequence with overlaps was gained via PCR using NEB Q5 polymerase with pGA-01-01 as template using primers `MagLOV_EcoFlex_FWD` and `MagLOV_EcoFlex_REV`. Both plasmids were combined in a subsequent Level 2 EcoFlex reaction with the pTU2-a backbone, leading to pVS-02-04_mCherry+MagLOV. This plasmid was transformed into *E. coli* MG1655 for mother machine experiments. The predicted translation rate is 250.

*pGA-01-47* The sequence for MagLOV 2 was codon optimised and the RBS designed to maximise consitutive expression (c.f. *pRSET-AsLOV_R2, pRSET-MagLOV, pRSET-MagLOV-2, pRSET-MagLOV-2_R11-f* which are inducible) using the RBS Calculator and CDS Calculator tools of [5]. The sequence was synthesized as a gene fragment (Twist Bioscience) and assembled into a level 1 plasmid as above, again using the J23119 promoter. This plasmid was transformed into *E. coli* MG1655 for multiplexing experiments.

*pRH-01-17* Same as pGA-01-47 but for AsLOV2 R5.

For pVS-02-04, an unoptimised codon sequence and RBS were used resulting in a predicted translation rate of 261 [5]. For pGA-01-47 and pRH-01-17 the optimised codon sequence and RBS yielded a predicted translation rate of $10^6$, a $20\times$ increase in predicted rate over the pRSET strains and a $4000\times$ increase in predicted translation over pVS-02-04. Note these predictions also do not account for the dual expression in pVS-02-04 of mCherry.

#### S1.5 Microfluidic Characterisation

Microfluidic experiments were performed in a "mother machine" microfluidic chip [6]. Microfluidic devices were cast in PDMS (Dow Corning Sylgard 184 kit, ConRo - 1673921) from a silicon wafer template (gift from Stephan Uphoff). Monomer and curing agent were combined 1:10, degassed in a vacuum to remove bubbles and cured at 65C for 2 hours. PDMS devices were cut out with a scalpel and inlet/outlet holes punched with a 0.75mm biopsy punch (Labtech - 52-004908). To prepare for bonding, PDMS devices were washed in isopropanol and dried with a manual dust blower. Precision cover glasses (1.5H thickness, Thorlabs - CG15KH1) were prepared for bonding by 5 minutes sonication in acetone, followed by 5 minutes sonication in isopropanol and drying with a manual dust blower. Clean PDMS devices and cover glasses were treated in an HPT-100 Henniker Plasma plasma oven (65% power, 0.3mbar, 30 seconds). Subsequent bonding of PDMS to cover glass was performed on a hot plate at 80C, followed by 1 hour baking at 95C.

Cultures used in mother machine experiments were incubated overnight at 37C 250rpm in M9 medium (Formedium - MMS0102) with the corresponding antibiotic. Three hours before loading, cultures were diluted 1:100 in fresh M9 medium containing 0.2g/L Pluronic F-127 (Sigma Aldrich - P2443-250G) and antibiotic. Thirty minutes before loading, the chip was flushed with 10ul 2g/L Pluronic F-127 using a P200 pipette with a gel loading pipette tip (Bio Rad - 12021138). After three hours growth (reaching OD600 0.2∼0.4), cell cultures were OD matched, and mixtures were prepared. For lock-in experiments (mixture of EGFP and MagLOV) the cells in the mixtures had different antibiotic resistance, hence no antibiotic was used after this point. The cells then were loaded into the chip in the same manner as above. The loaded chip was then centrifuged twice for 3 minutes, 1250RCF at a 10 degree angle to force cells into the trenches on both sides of the chip. Once connected to microfluidics, cells were supplied with M9 medium containing 0.2g/L Pluronic F-127 at a flow rate of $15 - 20\,\mu L/min$ (controlled by an Elveflow OB1 Mk3+ Flow Controller, Darwin Microfluidic - LVF-OB1). Cells were maintained at a constant temperature of 37C during the experiment.

#### S1.6 Microscopy Hardware

All images were acquired using a custom built microscopy platform [Fig. S1] based on the Rapid Automated Modular Microscope System (RAMM) from Applied Scientific Instrumentation (ASI). The configuration includes a Nikon PLAN APO $\lambda$D 40x/0.95 N.A. objective (part no. MRD70470), a Teledyne Kinetix sCMOS Camera, a five-band fluorescence filter set (405/445/514/561/640nm Quinta Band Set, Chroma part no. 89904), and Nikon Ti-2 200mm tube lens (36mm clear aperture), resulting in a pixel size of $0.1625\,\mu m \times 0.1625\,\mu m$ and a field of view (for the $3200 \times 3200$ pixel resolution camera) of 0.52 mm $\times$ 0.52 mm. The microscopy hardware was driven by a custom-built synchronisation electronics board (built around the Teensy 4.1 ARM Cortex-M7 microcontroller) and a custom-built LED driver system which can provide up to 70W of power to each of up to eight LEDs channels with hardware-based feedback control of LED current or optical output power. The system was configured with five LEDs with wavelength centres at 385/450/515/565/645nm (Luminus Devices part numbers CBM-50X-UV-Y31-FA380-22, SBR-70-B-R75-KG301, PT-39-G-L51, CBT-90-CG-L11-G100, and PT-39-DR-L51-BD100 respectively). LED light was homongenised using a two-layer micro-lens array and 4f relay system to flatten excitation light intensity across the field-of-view (approximately 5% variation from centre to corner extremity), and projected through a Texas Instruments Digital Micromirror Device (DMD) system (part numbers DLPLCRC900DEVM and DLPLCR67EVM). The microscopy platform was integrated and automated using custom driver software and operating system/GUI written in Python (control/processing computer) and C++ (microcontroller).

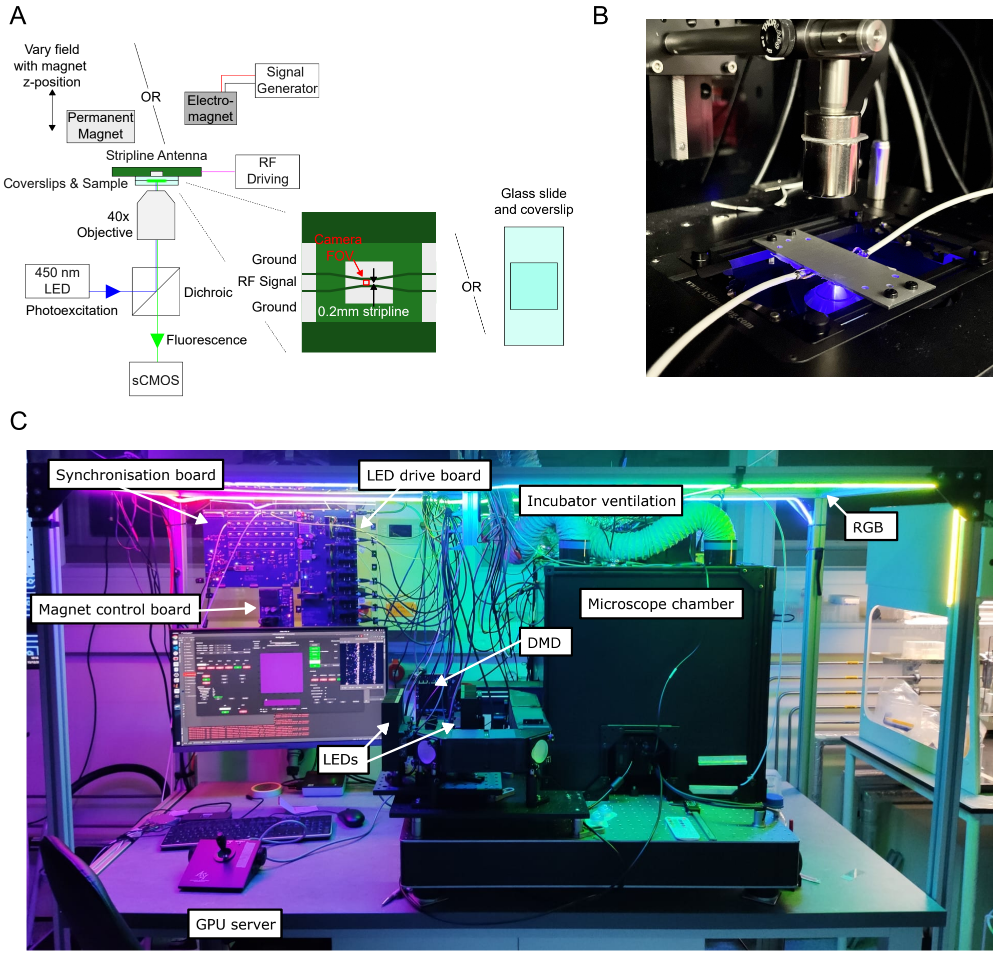


**Fig. S1**: **A.** Schematic of the inverted epi-fluorescent microscope with integrated RF components. The sample is either imaged near to a printed circuit board (PCB) stripline antenna as shown for ODMR and MFE experiments (to ensure consistent background light when comparing MFE and RFE), or on a glass-slide for multiplexing MFE experiments. A static magnetic field is provided either by a permanent magnet for ODMR (as in B.) or by an electromagnet. **B.** Photograph of the microscope stage in configuration for ODMR measurements. The inverted PCB is shown, with coaxial cables connected to the stripline, and permanent magnet above this in a raised position. **C.** Photograph of the custom built microscopy platform (as in A.) used for MFE/ODMR characterisation and microfluidic experiments.

#### S1.7 Acquisition Sequencing

Signal triggering and LED control is performed by the electronics board described above, and depicted for each experiment in Fig. S2. The microcontroller is programmed to generate signals to trigger single image acquisitions, as well as to step the RF frequency (for ODMR experiments) or set the magnetic field strength (for MFE experiments). It also controls the LEDs. Therefore the RF, LED, electromagnet field and camera state are all synchronised to within $\sim 10\,\mu s$. While the acquisition is ongoing, controlled by the microcontroller, the computer asynchronously receives images from the camera through a high-speed PCIe interface, approximately at the rate they are acquired.

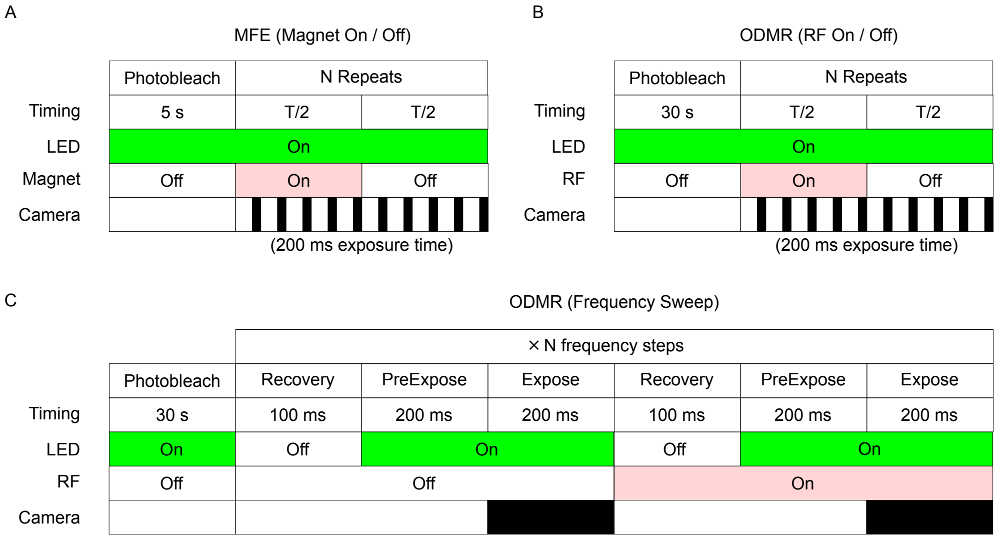


**Fig. S2**: **A.** Trigger sequence for magnet on/off imaging, as in Fig. S1, Main Text Figs. 1, 2, 4 and 6. For the camera sequence, filled-black indicates exposing. **B.** Trigger sequence for ODMR as in Main Text Fig. 2. **C.** Trigger sequence for ODMR as in Main Text Fig. 1 D and E.

#### S1.8 Magnetic Field Generation and Measurement

Widefield microscopy magnetic field effect experiments were performed on the microscope described in S1.6, with the addition of an iron core electromagnet (KK-PO50/27) driven by a custom-built bipolar current supply printed circuit board (PCB). A schematic is shown in Fig. S3 A, and the assembled board in Fig. S3 B. An operational amplifier (OpAmp, OPA549T) is used to amplify an analog control signal generated by the synchronisation board. The signal input can either be an arbitrary waveform generated by the digital-to-analogue converter (DAC), or it can be modulated by the IC Switch at high speed. This allows either a simple square wave by setting DAC1,2 to constant values and switching between them, or other lock-in modulation schemes where both DAC1,2 carry a time-varying signal that is switched between. The input signal ranges between $0-3.3$ V where $3.3/2$ V is approximately the zero-current point; this can be calibrated using the current sense resistor. Current is controlled using a current amplifier circuit (with 25mOhm $\pm 20$ ppm/C sense resistor, sense OpAmp

INA296) feedback loop with an (analog) error integrator to account for steady-state error. The ground plane is split such that large currents generated by the ±24 V supplies (top right of Fig. S3 B) are isolated from the digital and analogue low-power signals that perform sensing and control (lower half of Fig. S3 B).

Magnetic fields generated by the electromagnet and the permanent magnet were measured in the $z$-axis (Phidget Hall Effect Magnetic Sensor 1108_0). For data reported in the main text, the electromagnet was switched between zero and a fixed positive current, however we performed similar experiments alternating between equal magnitude positive and negative currents and saw no difference in results.

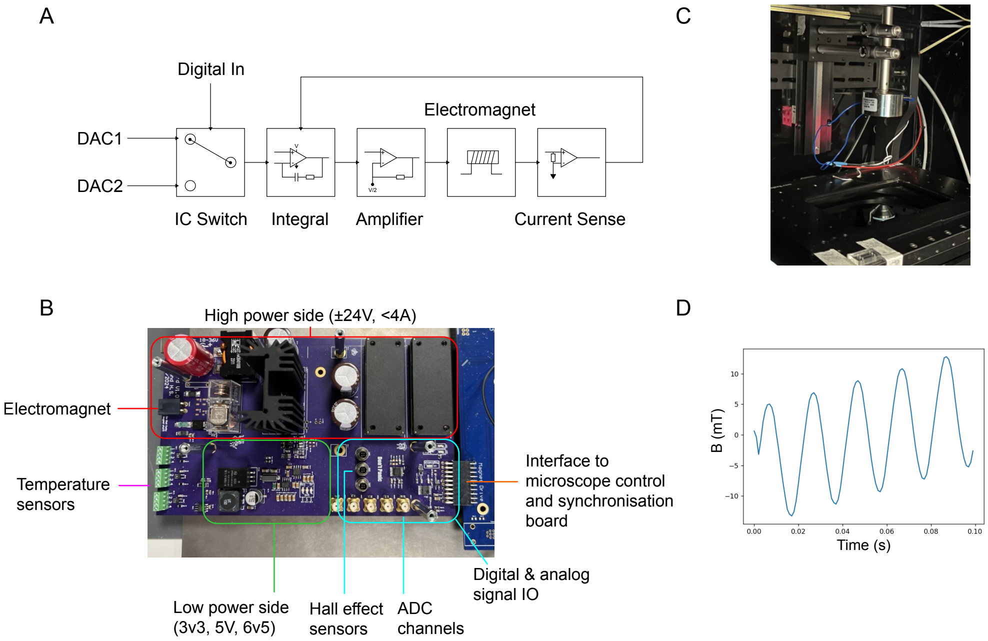


**Fig. S3**: **A.** Schematic of the electromagnet current supply electronics printed circuit board (PCB). **B.** Assembled electromagnet supply PCB. **C.** Photograph of the microscope stage with electromagnet (in raised position). **D.** Hall sensor measurement of the electromagnet, with the supply programmed to generate a current with 20ms period sin wave and linear ramp, demonstrating fast and accurate current control and magnetic field readout.

#### S1.9 Genetic Controls

We recorded an ODMR spectrum from MagLOV and compared with a negative control [Fig. S4 A], revealing an ODMR signal is present only when the MagLOV protein is expressed. We grew *E. coli* expressing MagLOV, and *E. coli* transformed with the same plasmid devoid of the MagLOV protein coding sequence (negative control), scraped samples of each and placed cell conglomerates next to each other

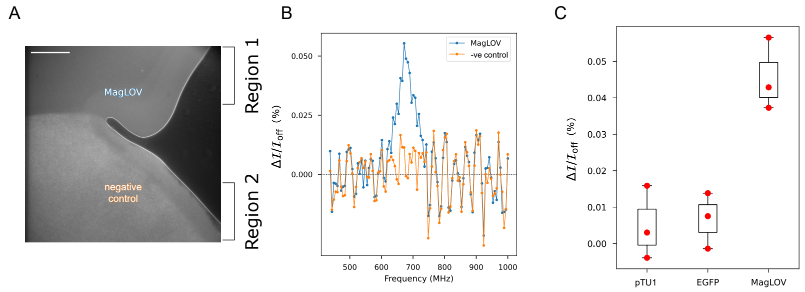


**Fig. S4**: **A.** Single field of view with MagLOV and negative control cells present as used in panels B/C. Scale bar is 100 µm. LED illumination is 450 nm, 1.5 W/cm$^2$. **B.** ODMR trace of bulk MagLOV cells and negative control cells in the same field of view, found by integrating horizontal regions 1 and 2 of panel A. Note that in this data the RF antenna was *not* calibrated to ensure a uniform power delivery across frequency (as explained in Fig. S7); we hypothesise that the aligned peaks observable in both the MagLOV and negative control samples are attributable to changes in delivered RF power from the non-uniform antenna response leading to secondary effects of RF power on the imaging setup. **C.** ODMR of bulk cells (MagLOV 2 fast) (similar to A, but only one type of cell), performed in biological triplicates at the same magnetic field $B_0 \approx 24.3\,\mathrm{mT}$ to compare peak ODMR contrast measured at the resonant frequency 680 MHz. The box extends from the first quartile to the third quartile of the data, the whiskers extend from the box to the farthest data point lying within 1.5x the inter-quartile range from the box.

in the same field of view above the stripline [Fig. S4 A]. Integrating the brightness of the signal over the labelled regions of the image yields the MagLOV curve and negative control curves in [Fig. S4 B] (note that negative control cells are visible due to autofluorescence [7]). In the fluorescence signal from both cell types, we see non-magnetic resonance effects, hypothesised to arise from resonance effects inherent in the experimental hardware and non-uniformity in RF amplifier output (see Fig. S7). However, the MagLOV protein fluorescence *also* exhibits an ODMR signal at the expected ESR frequency of $\omega_{RF} = \bar{\gamma}_e B_0 = 684\,\mathrm{MHz}$ for $B_0 = 24.4\,\mathrm{mT}$. Furthermore, we imaged cell conglomerates transformed with an empty plasmid (negative control), MagLOV plasmid (same as empty plus MagLOV), and an identical plasmid where MagLOV is replaced with EGFP [8] (which is much brighter than MagLOV) in biological triplicates, and only observed an ODMR resonance in cells expressing MagLOV [Fig. S4 C].

Compared to the single cell data (Fig. 2), the bulk measurement contrast is reduced (for example we saw an ODMR contrast of $\sim 0.05\%$ in Fig. S4 B) compared to single cells. This is likely because of increased background fluorescence due to cell debris, and strongly auto-fluorescent LB-agarose that may be collected when picking the colony from the agar-pad.

#### S1.10 RF Hardware

The RF chain is shown in Fig. S5 A. An RF signal generator (Windfreak SynthNV Pro) is programmed to generate signals at $P_{out} = 0 - 2$ dBm. The signal generator is trigger controlled (by the microscope synchronisation board) to step between each frequency. The output of the radio is attenuated by 15 dB (Mouser 549-CATTEN-0150), then passed through an RF switch (Mini-Circuits ZYSWA-2-50DR+) that is controlled by another digital signal. The signal is then amplified by $\sim$ 50 dB (Mini-Circuits ZHL-20W-13S+), and fed into the stripline antenna PCB. The output of the antenna is attenuated by 40 dB (Mouser 523-2082-6525-18-40), and fed back into WindFreak as a diagnostic that radio is on and circuit is connected (the WindFreak measures total power input, but is not spectrally resolved).

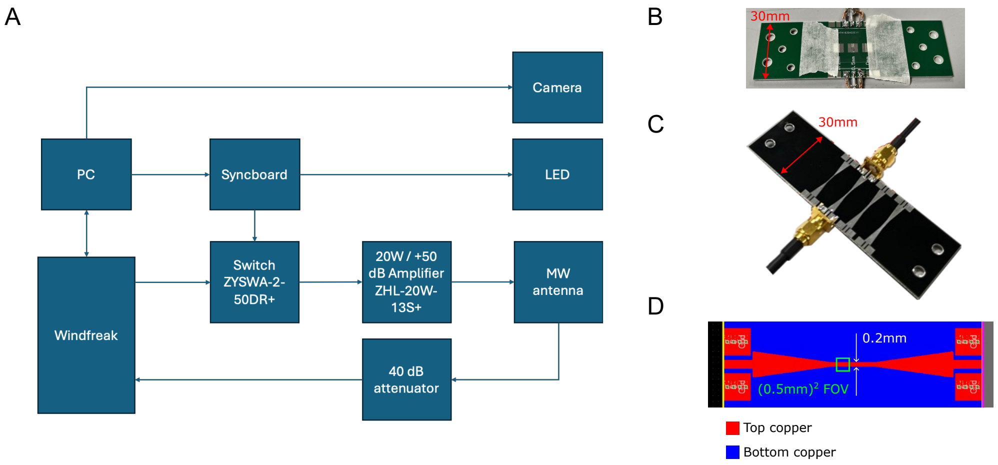


**Fig. S5**: **A.** Components used in the widefield ODMR setup. **B.** First version of stripline antenna PCB, with glass coverslides attached. **C.** Redesigned antenna with multiple stripline widths shown. The 0.2 mm stripline was selected as it provided a good $B_1$ field strength and cells across the entire FOV and exhibited clear ODMR signals. **D.** PCB layout of the redesigned antenna, cropped to the 0.2 mm stripline. *Not shown:* a piece of black tape was placed over the antenna (under the cover slips) to remove reflections from the antenna metal. ODMR measurements were performed with and without the tape with no significant difference apart from image quality (i.e. same line-shapes observed, but lower contrast without the tape due to the contribution of reflected excitation light).

#### S1.11 Determining Protein Concentrations

##### S1.11.1 Purified Protein

Purified stocks of MagLOV 2 and MagLOV 2 fast were prepared in PBS buffer with 30% glycerol, and stored at −80 °C. Following the method Wingen et al. [9] used to determine the extinction coefficient of various LOV-based fluorescent proteins, we estimated the concentration the purified protein samples, by first heating them to 95 °C for 5 minutes to dissociate the FMN from the protein (we assume that one FMN corresponds to one protein, since free FMN should be removed by the protein purification process). The concentration of flavin (and hence protein) was then determined by performing a serial dilution and measuring $A_{450\ \mathrm{nm}}$, using the free FMN extinction coefficient $\epsilon_{\mathrm{FMN}} = 12,200\ \mathrm{M}^{-1}\mathrm{cm}^{-1}$ [10].

##### S1.11.2 Intracellular Protein Concentration

Intracellular protein concentration of MagLOV 2 in T7 Express strains (as used in Fig. 1 and 2) was estimated using the FPCountR method [11], approximating the extinction coefficient of MagLOV 2 at 14200 $\mathrm{M}^{-1}\ \mathrm{cm}^{-1}$ in line with other fluorescent LOV-binding proteins [9]. We found an intracellular concentration of $10 \pm 2\ \mu$M across 3 biological replicates $\times$ 3 technical replicates.

#### S1.12 Magnetic Sensitivity

We estimated the sensitivity of MagLOV variants by calculating the static magnetic sensitivity in a CW ODMR measurement in photon shot noise limit according to

$$\eta_0 = \frac{\mathcal{P}_{\mathcal{F}}}{\gamma_e} \frac{\Delta\nu}{C\sqrt{R_0}}$$

where $\mathcal{P}_{\mathcal{F}}$ captures the peak shape (either $\sim$ 0.70 for Gaussian or $\sim$ 0.77 for Lorentzian), $\Delta\nu$ is the line width, $C$ is the contrast, $\gamma_e = 28\,\mathrm{MHz/mT}$ is the electron gyromagnetic ratio, and $R_0$ is the photon count rate [12].

For purified MagLOV 2, we find that $\eta_0 = 0.4\ \mu\mathrm{T\ Hz}^{-1/2}$ when integrating over the entire field of view. As a more meaningful figure of merit and to facilitate comparison with other works, we estimate the concentration normalised sensitivity by multiplying the sensitivity by the number of molecules present in the sample,

$$\eta = \eta_0 \times \sqrt{cV}$$

where $c$ is the concentration and $V$ is the volume of the sample. Our useful microscope field of view is $0.5 \times 0.5\ \mathrm{mm}^2$, and we measured the distance from coverslip to coverslip as 2.5 $\mu$m from sharp image plane to sharp image plane at the edge of the liquid drop (indicating the liquid-glass interface). In this case we find the purified protein solution has a room temperature sensitivity of $\eta = 53\ \mathrm{fT\ mol}^{1/2}\ \mathrm{Hz}^{-1/2}$ [Fig. S6 and Table. S1]. Interestingly the purified protein has a significantly decreased ODMR contrast compared to *in-vivo* measurements (e.g. Fig. 1), suggesting the cellular environment has a large impact on the spin-radical dynamics.

It is also interesting to estimate the magnetic sensitivity of a single cell. For single cells expressing MagLOV 2, with illumination intensity of 800 $\mathrm{mW/cm}^2$ at 450 nm as in Main Text Fig. 1 D, we estimate the sensitivity by integrating the light

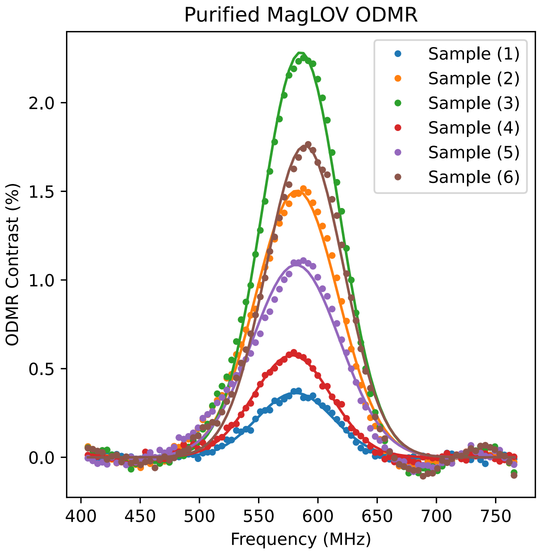


| Sample | MagLOV 2 1 | 2 | 3 | MagLOV 2 fast 4 | 5 | 6 |
|---|---|---|---|---|---|---|
| Concentration ($uM$) | 2.55 | 25.50 | 25.50 | 1.88 | 18.80 | 18.80 |
| LED Intensity (mW/cm$^2$) | 1477 | 351 | 1477 | 1477 | 351 | 1477 |
| FWHM (MHz) | 75.78 | 82.08 | 77.09 | 77.31 | 87.07 | 77.43 |
| Contrast (%) | 0.38 | 1.52 | 2.25 | 0.59 | 1.11 | 1.76 |
| Photon Count Rate (Hz) | 2.33e+10 | 1.90e+10 | 4.52e+10 | 2.66e+10 | 2.63e+10 | 4.11e+10 |
| $\eta_0$ ($\mu$T Hz$^{-1/2}$) | 3.31 | 0.98 | 0.40 | 2.00 | 1.21 | 0.54 |
| $\eta$ (fT mol$^{1/2}$ Hz$^{-1/2}$) | 137.52 | 128.90 | 52.82 | 71.30 | 136.31 | 60.98 |

Fig. S6 & Table S1: Purified protein ODMR measurements. Fit lines are Gaussian ($\mathcal{P} = 0.7$ in $\eta$ calculation).

collected by pixels that cover a single cell. We find an average integrated cell brightness of $5.75 \times 10^5$ Hz, 10% contrast, 80 MHz linewidth, giving a per cell sensitivity of $\eta_0 = 26$ $\mu$T Hz$^{-1/2}$. We also estimated the intracellular protein concentration of MagLOV 2 at 10 $\mu$M (see Section S1.11.2). Taking the typical cell volume as 1 $\mu$m$^3$ [13], we find a per cell molar sensitivity of $\eta = 2.6$ fT mol$^{1/2}$ Hz$^{1/2}$.

From the MARY data [Fig. S12] collected for MagLOV 2 fast, we can also estimate the sensitivity to $B_0$ detected via MFE. In this case, we have $\eta_0 = \frac{1}{\delta MFE/\delta B_0}\frac{1}{\sqrt{R}}$, where $\delta MFE/\delta B_0 \approx 4.7$ T$^{-1}$. We find that $\eta_0 = 52.5$ $\mu$T Hz$^{-1/2}$, and $\eta = 1.70$ nT mol$^{1/2}$ Hz$^{-1/2}$ under the conditions measured (200 $\mu$L volume, 100 mW/cm$^2$). This change in sensitivity compared to ODMR is as a result of the much larger line-width of the MARY curve compared to the ODMR; the MARY line-width of 26 mT [Fig. S12] corresponds to a 730 MHz shift, roughly 10× the ODMR line-width (also see the simulation work in Section S1.18 for further discussion of this).

Next, we compare the MagLOV sensitivity with some relevant systems from the literature, namely other protein and organic molecules, see Table S2. It is important to note that the sensitivity $\eta_0$ as given above is context dependent, for instance the larger the sample size measured the larger the photon count rate will be, thus reducing (improving) the calculated sensitivity. Therefore it is difficult in most cases to make direct comparisons, except where other works have reported the photon count, and/or reported or normalised by the sample volume. In general, holding other parameters constant, increasing contrast and decreasing line-width both lead to improved sensitivity.

With regards to non-protein based spin sensors that require introduction to the cell, nanodiamonds stand out as having superior sensitivity and low photobleaching, and in cells have sensitivities reported broadly in the range $2.6 - 5.3\,\mu T/\sqrt{Hz}$ with *in-vitro* samples at the lower end and *in-vivo* at the higher [14–17]. While MagLOV is unlikely to reach these sensitivities given the large linewidth of $\sim 80$ MHz (though comparatively similar brightness and contrast), the advantage of a protein sensor is that it is not merely biocompatible, but produced by the cell itself, eliminating much of the complications required to introduce NV nanodiamonds (or alternative molecules) into the cell, and unlocking a broad range of novel application possibilities. For instance, MagLOV could act as a genetic reporter as a fusion protein, whereas any foreign system would first require introduction to the cell followed by attachment via a biological binder.

| System | Sample Type | Temperature | Contrast (%) | Line-width (MHz) | Reference |
|---|---|---|---|---|---|
| DmCry | purified | RT | 2 | ∼ 100 | [18] |
| EYFP | purified | RT | ∼3 | ∼55 | [19] |
| EYFP | frozen HEK 293T cells | 175 K | Not Reported | ∼55 | [19] |
| m-Scarlet+FMN | purified | 294.65 K | ∼7.5 (max) | ∼100 | [20] |
| m-Scarlet+FMN | live nematode | 294.65 K | ∼1.5 (max) | ∼100 | [20] |
| MagLOV 2 | purified | RT | 2.25 | 77.1 | Fig S6 Sample 3 |
| MagLOV 2 | live *E. Coli* cell | RT | ∼10 | 80 | Fig. 1 main text |
| MagLOV 2 fast | purified | RT | 1.76 | 77.4 | Fig S6 Sample 6 |
| NV nanodiamonds | live *C. elegans* | RT | 12 | 22 | [21] |

**Table S2**: Comparison of recently reported contrast ODMR contrasts for similar protein systems, and an exemplar nanodiamond measurement. RT: Room Temperature

#### S1.13 Stripline Antenna Design

The stripline printed circuit board (PCB) antenna was designed in-house through several iterations of design and testing (i.e. a PCB directed evolution protocol selecting for response magnitude) that varied geometry and layout to optimise RF power delivery within the sample area (measured by a small loop RF antenna), and manufactured by PCBWay [Fig. S5 B-D]. The final antenna design is approximately 0.035mm in thick (corresponding to standard 1oz PCB copper traces) and 0.2 mm in width, which tapers to solder pads at the edge, following a similar geometry to that proposed by [22]. The PCB structural material is 1.5 mm thick aluminium (to act as a heat sink) and the surface finish is HASL with lead. A piece of black electrical tape was placed over the antenna / underneath the bottom cover for imaging to remove LED reflected light (not shown in Fig. S5 B,C).

##### S1.13.1 Stripline Antenna Response

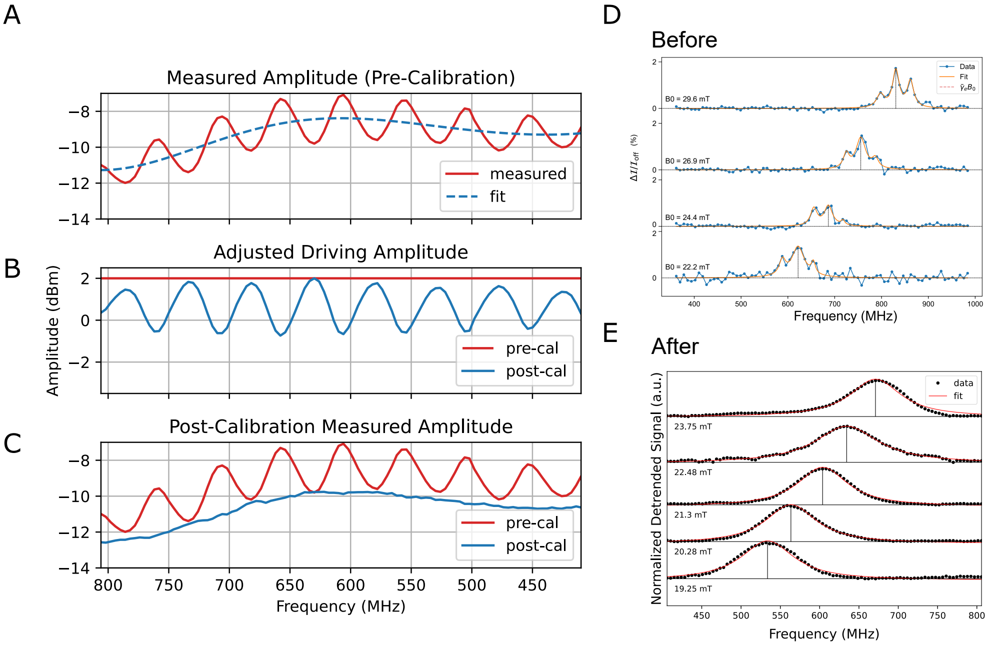


**Fig. S7**: **A.** Power meter measurements performed by the radio signal generator during a frequency sweep with the stripline antenna connected. Red solid line is the total measured power at each driving frequency. Blue dashed line is a smooth fit to the red line, which is the target power output. We used this smooth fit rather than a flat line as the target, as otherwise the final power output would be severely limited having to compensate for a 5dB difference from maximum to minimum of the measured spectrum. **B.** The driving power curve. Red line shows pre calibration (which results in the measured line in A.). Blue line shows the new power curve designed to achieve the dashed blue line in A. **C.** Post calibration, the power curve (blue) is significantly flatter than the original measured curve (red). This removes the 3 peak structure exhibited in D. **D.** ODMR spectra at various $B_0$ fields prior to calibration exhibiting resonance artefacts. **E.** ODMR spectra at various $B_0$ fields with calibration, no longer exhibiting artefacts.

For widefield ODMR imaging, as the sample is in the near field, the $B_1$ field strength can be estimated using the Biot-Savart law. From Fig. S7, with the radio driving at 0 dBm at 600 MHz, followed by −15 dB attenuation and +50 dB amplification, and an additional −40 dB attenuation prior to the measured amplitude of −10 dBm, we find that the system has 5 dB of losses meaning the power through the antenna is $\sim 30$ dBm. Using $B_1 = \frac{\mu_0 I}{2\pi z}$, where $z = 0.17$ mm (coverslip thickness), current $I = \sqrt{P/Z}$, $P = 30$ dBm and $Z = 50\ \Omega$ we find $B_1 \approx 0.167$ mT. Experiments shown in the main text had the radio set to +2 dBm (so $P = 32$ dBm), giving $B_1 \approx 0.2$ mT.

Initially, we assumed the peak-like structure in Fig. S7 was a result of FMN hyperfine lines. However, further investigation revealed strong resonance structures in the antenna, which coincide with the peaks in Fig. S7 D. This antenna [Fig. S5 B] also exhibited significant line losses, meaning the correction scheme described in Fig. S7 would not have been possible, as the corrected RF power would be too low. This motivated the iterative re-design of the antenna described above. The key improvements were the addition of edge connectors, which improved the matching to the coaxial cable, and the removal of the antenna-side ground plane (i.e. switching from a co-planar to a micro-strip design) [Fig. S5 C,D]. With these changes, we measured an 18 dB increase in $B_1$ field at the antenna surface (using a pickup loop and Rigol DSA875 spectrum analyzer), yielding the red curves in Fig. S7 A,C which was then corrected to the blue curve in Fig. S7 C. This calibration procedure was performed prior to every ODMR experiment to ensure uniformity of RF power delivery across the scanned frequency range.

#### S1.14 Sample Preparation

Plasmids expressing EGFP, MFP variants, or negative control plasmids expressing only antibiotic selection markers were used to transform New England Biolabs T7 Express Competent *E. coli* cells (C2566H) via heatshock at 42 °C for 45 seconds. Colonies were grown overnight at 37 °C on Agar plates, then a single colony was picked, re-suspended in 1 mL of LB media, spread onto a plate, and grown at 37 °C for 24 hrs followed by 24 hrs at room temperature to form a lawn of cells.

##### S1.14.1 Bulk Widefield Measurements

For ODMR, cells were then scraped from an agar plate directly onto a glass cover slip (SLS MIC2162). A second cover was placed on top and then both placed atop the microwave antenna stripline (Fig. S5 C) and secured with tape. For MFE, cells were scraped onto a glass slide (VWR Superfrost) and a glass cover was placed directly on top.

##### S1.14.2 Single-Cell Resolved Widefield Measurements

For ODMR, MagLOV *E. coli* cells spread from a single clone were scraped off an agar plate and resuspended in PBS buffer, then a $\sim 1\,\mu$L droplet was sandwiched between two glass covers taped to a stripline antenna, creating a monolayer of roughly 1000 individually distinguishable cells per $0.5 \times 0.5$ mm$^2$ field of view (FOV). For MFE, the same procedure was performed but onto a glass slide with a cover placed on top.

##### S1.14.3 Multiplexing

For the data in Main Text Fig. 4 B-C, MagLOV 2-R11-f and AsLOV2-R2 were resuspended in PBS and made up to equal concentration as measured by optical density (OD600). Individually, and as a $1:1$ mixture, monolayers of cells were sandwiched between a glass slide and cover for MFE imaging.

#### S1.15 Widefield Imaging Protocols

All measurements performed at room temperature, apart from microfluidic experiments where temperature is held at $\sim 37°$C to promote cell growth.

##### S1.15.1 Widefield Light Source Calibration

| **Figure** | **Power Density** |
|---|---|
| Fig. 1 C,D | 800 mW/cm$^2$ |
| Fig. 1 E | 2.8 W/cm$^2$ |
| Fig. 2 | 800 mW/cm$^2$ |
| Fig. 4 B, C | 800 mW/cm$^2$ |
| Fig. 4 F | 1.5 W/cm$^2$ |
| Fig. 5 B | 800 mW/cm$^2$ |
| Fig. 6 | 280 mW/cm$^2$ |

**Table S3**: Power densities used in main text figures.

LED power calibration was performed by using the microscope LED system DMD to project a square of uniform illumination that would cover a known number of (camera) pixels onto a ThorLabs PM100D power meter positioned at the sample position. 450 nm LED power levels are given in Table S3.

##### S1.15.2 Magnetic Field Effect Measurements

MFE curves are acquired under continuous LED illumination, with an electromagnet switched on and off at a fixed period $T_{\mathrm{magnet}}$. Photobleaching effects dominate the fluorescence curve at the beginning. Camera exposure time and LED intensity were chosen to balance rate of photobleaching, signal-to-noise, total acquisition time, and the dynamics of the MFE (if LED intensity is increased too high, images can not be taken fast enough to see the dynamics due to a frame-rate limitation of at least 10 ms between each frame).

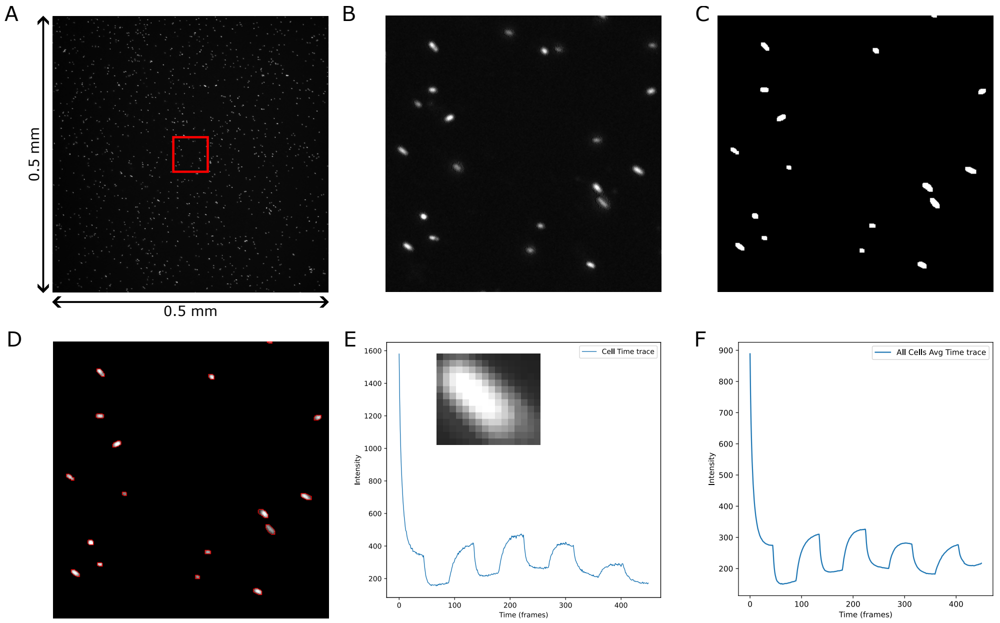


**Fig. S8**: **A.** A single field of view (FOV) is shown, with a crop region for the purposes of this figure (normally the entire FOV is processed). **B.** The cropped region, showing single *E. Coli* cells expressing fluorescent MagLOV. **C.** The image is masked, using a brightness threshold followed by opening. **D.** From the mask, cells are identified (methodology described in the text). **E.** A single cell is shown, and it's fluorescent time trace as a result of applying the mask in E. to the full time series cropped to that cell. Frames are collected every 200ms as in Fig. S2. **F.** The overall average trace is shown, by averaging over all the individual cell time traces.

**Time-Trace Extraction.** For images of single cell monolayers, cell regions were segmented using brightness thresholding and morphological operations. These image processing steps were performed using functions (`morphologyEx` (with `MORPH_OPEN` followed by `MORPH_CLOSE`) and `findContours`) from the Open Source Computer Vision Library (OpenCV, version 4.10.0).[23] as illustrated in Fig. S8. Opening and closing are techniques to remove noise (i.e. bright (dark) pixels smaller than the size of a cell against a background of by dark (bright) pixels). The fluorescence time trace was then computed for each cell by averaging pixel intensities within cell masks, across each image.

**Baseline Subtraction and Fitting.** The intensity time trace as extracted above exhibits two modes of baseline that we remove to extract the modulated signal. Firstly, upon illumination there is an immediate and fast rate of photobleaching [Fig. S9 A]. This is common across all radical-pair based fluorescent proteins, for instance see [24] and is removed simply by discarding the first few seconds (in our case 10 seconds) of the acquisition before further processing. Secondly, there is a slow moving baseline

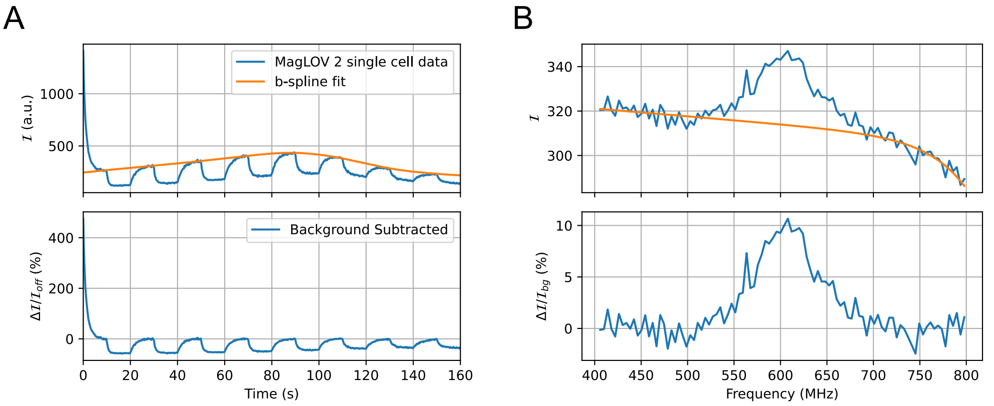


**Fig. S9**: **A.** Single cell MFE data (same data as in Fig. 1 C) before and after background subtraction. **B.** Single cell ODMR data (same data as in Fig. 1 D).

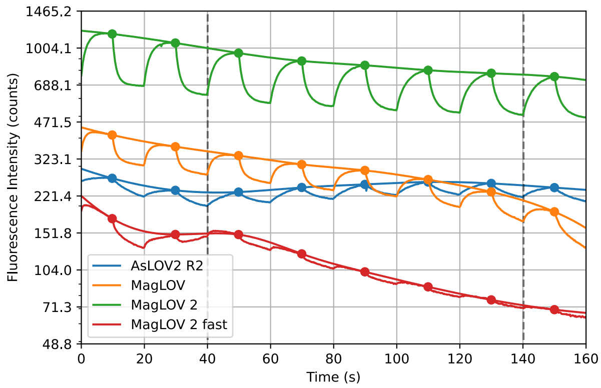


**Fig. S10**: Raw data for Fig. 2. Vertical dashed lines indicate the region in which periods were averaged.

drift. For MFE and RFE experiments we remove this baseline drift by fitting a b-spline (`make_interp_spline` in the SciPy package [25]) to the first point in the period before the magnet (or radio) is switched on. For lock-in MFE experiments [Fig. 4 F] we constructed the baseline by performing a moving average.

Following baseline subtraction, periods were averaged together (i.e. going from the time trace in Fig. S9 A to the single period in Main Text Fig. 2). Parameters were extracted by fitting an exponential model [26] (amplitude and growth rate)

$$y(t) = \begin{cases} a_1(1 - e^{-t/\tau})/(1 - e^{-T/2\tau}) + b_1, & 0 \leq t < T/2 \\ a_2\left[1 - (1 - e^{-(t-T/2)/\tau})/(1 - e^{-T/2\tau})\right] + b_2, & T/2 < t \leq T \end{cases} \tag{1}$$

where $\tau$ is the saturation timescale, $T$ is the period, and $a_i, b_i$ capture the MFE contrast and account for background offset. The distributions of these parameters were analysed across all cells as discussed in the main text.

##### S1.15.3 ODMR Measurements

To acquire ODMR spectra, a radio-frequency generator (Windfreak Synth NV Pro) generated frequencies in a linear frequency sweep while the cells were illuminated by a 450 nm LED.

To reduce the effect of photobleaching, a recovery time (during which LED and RF are off) was introduced between each frame (as in Fig. S2). For cell conglomerates (as in Fig. S4), an entire region of the image was integrated to produce the time-series intensity.

An exponential curve is fit to the tails of the signal which is treated as the "background", $y_{fit}$. The normalised ODMR signal is then computed as $(y/y_{fit} - 1) \times 100\%$ [Fig. S9 B].

#### S1.16 Microfluidic Image Processing

For mother machine experiments, we acquired one reference image at 450 nm excitation and one at 565 nm, followed by recording a timeseries of 400 images at intervals of 0.25 s (4 Hz) with excitation at 450 nm (200 ms exposure time, 1.5 W/cm$^2$ LED intensity). The 450 nm reference images were then processed using the DeLTA 2.0 image processing pipeline [27] to obtain masks for both growth trenches and individual cells, allowing each cell to be assigned a particular trench. Trenches that were out-of-focus or contained no cells/filamenting cells were discarded. The cell masks were then applied to the 450 nm timeseries to extract mean fluorescence values (fluorescence per cell area) for each time point and cell. This procedure was repeated for 5 fields of view.

To generate a "ground truth" for the measurement we mixed cells expressing pVS-02-04_mCherry+MagLOV with cells expressing pVS-01-03_EGFP. Masks created from images at 450nm were applied to the 565 nm reference image. A cell was considered to be marked if its mean fluorescence value at 565 nm was greater than the 99th percentile of the fluorescence values in the 565 nm reference image.

To distinguish between the two cell populations based on their MFE, single-cell-level timeseries were processed as follows. First, the timeseries were filtered using a moving-average filter of window size 40 samples, corresponding to a cutoff frequency of 0.05 Hz. The metric to distinguish between "MFP-positive" and "MFP-negative" cells was then obtained by multiplying the smoothed signal with a square wave of period 0.1 Hz (corresponding to the magnetic field modulation period) and taking mean value.

#### S1.17 Spectroscopic characterization

##### S1.17.1 Wavelength-resolved Bulk Magnetic Field Effects

Wavelength-resolved MFE measurements (Main Text Fig. 3 and Fig. S11) were performed in bulk using a home-built cuvette-based fluorescence spectrometer described in [28]. Briefly, a 450 nm laser diode is used to excite the sample, which is housed inside custom-built Helmholtz coils. The emission is collected through a lens pair and dispersed by a spectrograph (Andor Holospec) onto a CCD array (Andor iDus420).

Cells were resuspended in PBS buffer at OD $\sim 0.3$ and placed in a quartz fluorescence cuvette (Hellma). Similarly to the widefield measurements, a magnetic field was switched on and off with field strength 10 mT and a period of 20 s (10 s on, 10 s off). The sample was illuminated at 450 nm (Oxxius LBX-450), at roughly 1 kW/m$^2$, and the emission was filtered using a 458 nm longpass filter (RazorEdge ultrasteep).

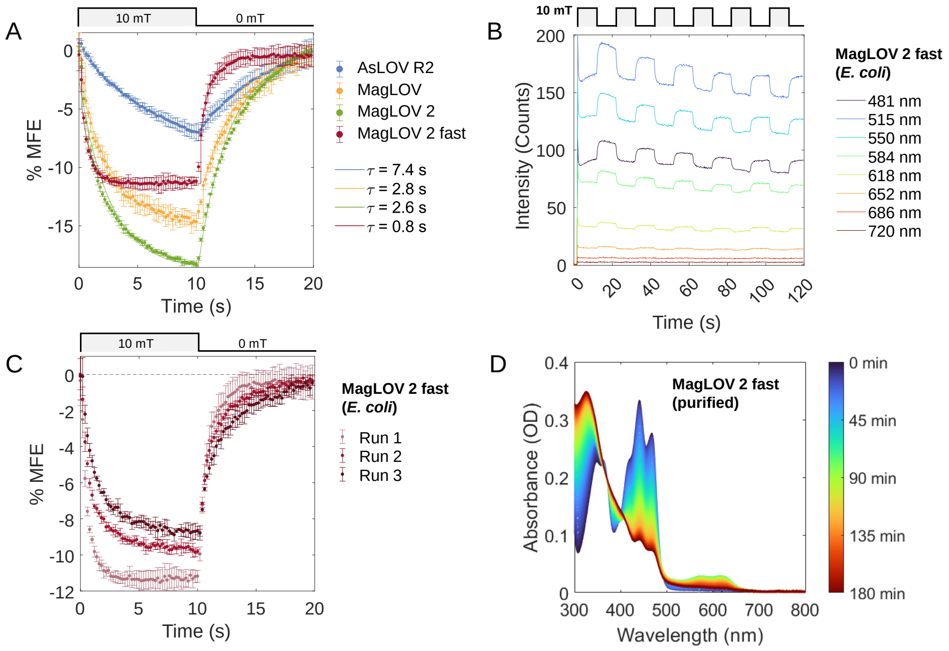


**Fig. S11**: **A**. Bulk measurements of each MFP variant in cells. The time constants ($\tau$) of the exponential fitted to the delayed MFE component are shown as solid lines. They are consistent with the widefield data [Fig. 2] given the samples are under different illumination conditions (here, light scattered through solution c.f. single cell monolayers) which we show to be a determinant of saturation rate (and hence contrast achieved within a given time) in Fig. S16. **B**. Representative time traces at a select number of emission wavelengths for MagLOV 2 fast. Note $B$ onset coincides with excitation. **C**. Comparison of three consecutive MFE experiments (each averaging 4 on-off steps), illustrating that some irreversible photo-conversion takes place over time, which reduces the MFE of a given sample. **D**. Change in absorption of (purified solution) MagLOV 2 fast under illumination of blue light. Colourbar indicates time after illumination onset.

##### S1.17.2 MARY Curve

To obtain the MARY (Magnetically Affected Reaction Yield) curve, the MFE of a purified MagLOV 2 fast solution was measured as a function of applied magnetic field strength, $B$, using the apparatus described above. Purified MagLOV 2 fast was diluted below $\sim$ 0.05 OD in PBS buffer (30% glycerol) and placed in a quartz fluorescence cuvette (Hellma). The sample was continuously excited at 450 nm (Oxxius LBX-450, $1\,\mathrm{kW/m^2}$), and the emission was filtered using a 458 nm longpass filter (RazorEdge ultrasteep). The magnitude and duration of the applied magnetic field and acquisition are controlled by software home-written (unfortunately) in LabVIEW. Eighteen

$B$ field points between $-20\,\mathrm{mT}$ and $20\,\mathrm{mT}$ are probed in a random order (50 repeats), with intermediate $0\,\mathrm{mT}$ measurements to minimise the effect of background instability such that data are collected as $(...,0_{n-1}, B_n, 0_{n+1},...)$. The MFE at $B_n$ is calculated relative to the mean of $0_{n-1}, 0_{n+1}$. A delay time ($2\,\mathrm{s}$) between field switching and spectrum recording (exposure time $200\,\mathrm{ms}$) is introduced to accommodate the MFE enhancement kinetics [24]. However, to minimise experimental time and sample degradation due to light exposure, the delay time is shorter than the saturation rate of the MFE. Consequently, the magnitude of the %MFE is comparatively smaller than in Figure S11.

The half-width at half maximum (conventionally known as the $B_{1/2}$) of the curve is extracted from a Lorentzian fit of the form $y = \frac{4Ax^2}{W^2+4x^2}$, where $A$ is the amplitude and $W$ is the full width at half maximum. For the data in [Figure S12], $A = (-3.0 \pm 0.3)\,\%\mathrm{MFE}$ and $W = (26 \pm 3)\,\mathrm{mT}$. Hence, $B_{1/2} \approx 13\,\mathrm{mT}$, which is comparable to the $B_{1/2}$ observed in flavomaquettes ($13-16$ mT), i.e. *de novo* designed flavoproteins, where the radical pair is formed between a riboflavin and tryptophan covalently attached to $\alpha$-helical bundles [29].

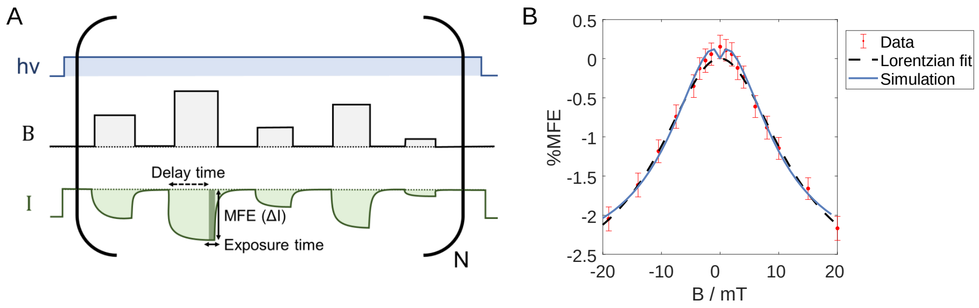


**Fig. S12**: **A**. Schematic diagram of the fluorescence MARY experiment. Under conditions of continuous illumination ($h\nu$, *blue*), a square wave magnetic field ($B$, *grey*) of different magnitudes is applied. This leads to a change in sample emission intensity ($I$, *green*). The MFE is quantified as $\%\mathrm{MFE} = \frac{I(B)-I(B=0)}{I(B=0)} \cdot 100\% = \frac{\Delta I}{I(B=0)} \cdot 100\%$. **B**. MARY curve for purified solution of MagLOV 2 fast (*red*). The half-width at half-maximum extracted from a Lorentzian fit (*black*), often denoted as $B_{1/2}$, is approximately $13\,\mathrm{mT}$ for both experiment and simulation. Simulation (*blue*) details in the following section.

##### S1.17.3 UV-Vis Absorption Spectroscopy

UV-Vis absorption spectra of the purified MagLOV 2 fast solution (Figures 3C, S11D, S13A) were recorded with an Agilent Cary 60 UV-Vis Spectrophotometer. The wavelength range from 300 nm to 800 nm was scanned every 0.2 minutes. Purified MagLOV 2 fast was diluted to OD$\sim$ 0.35 in PBS buffer, 30% glycerol, and placed in a quartz cuvette (Hellma, path length 1 cm). The sample was held at room temperature and continuously irradiated by a blue LED (XLamp XP-E2 LED, model XPEBBL-L1-0000-301, output ca. 10 mW at 450 nm) through the emission window, perpendicular to the UV-Vis probe beam.

##### S1.17.4 Emission, Excitation and Time-resolved Emission Spectra

Excitation spectra in Figures 3 C and S13 A were recorded using an Edinburgh Instruments FS5 Spectrofluorometer, using a Xenon lamp as an excitation source. Emission spectra [Figures 3 A] and time-resolved emission spectra [Figure S13C] were recorded on the same instrument, but using an HPL450 (450 nm) for excitation. Note that this instrument setup suffers from laser excitation artefacts which create the small peaks near 550 nm and 650 nm.

Steady-state emission and excitation spectra in the main text were recorded for a bulk cell suspension in PBS buffer at OD $\sim$ 0.3 (at 450 nm) and placed in a quartz fluorescence cuvette (Hellma). Time-resolved emission spectra [along with other emission data presented in Figure S13] were recorded on a solution of purified MagLOV 2 fast, diluted to $\sim$ 0.05 OD in PBS buffer (30% glycerol) and placed in a quartz fluorescence cuvette (Hellma).

Time-correlated single photon counting (TCSPC) curves (Figure S13B,C) were recorded with an HPL450 (10 MHz repetition rate) and a polariser on emission set at 55 degrees. Global reconvolution fitting was performed with the commercial Edinburgh Instruments software FAST. The hence determined fluorescence lifetime of MagLOV 2 fast in purified solution is 3.2 ns. Assuming the radiative lifetime of MagLOV-bound FMN is 16 ns, as in AsLOV WT (cf. 18 ns for free FMN) [30], the fluorescence quantum yield can be estimated as $\Phi_F \approx 3.2/16 = 0.2 \equiv 20\%$, which is comparable to other LOV domains [30–32]. Note that the radiative lifetime is an estimate, as the mutations introduced relative to the WT possibly alter the chromophore environment and non-radiative rates (intersystem crossing, electron transfer).

For the application to signal multiplexing described in the main text, both high fluorescence quantum yield and a substantial MFE are beneficial. In comparison to GFP derivatives (quantum efficiency $\Phi_F > 0.6$ [33]), MagLOV has a reduced $\Phi_F \approx 0.2$ [Figure S13]. Because fluorescence competes with radical-pair formation pathways such as intersystem crossing and subsequent electron transfer, efficient generation of the SCRP inevitably suppresses $\Phi_F$, creating a trade-off between MFE magnitude and brightness. However, in comparison to cryptochrome (singlet yield $\phi_s \sim 0.003$ (0.3%) or even below 0.1% [34, 35], with fluorescence-detected MFEs of up to 1% at 16 mT [26, 36]), MagLOV offers significantly improved brightness and MFE contrast at ca. $\sim$ 10% under similar conditions [Figure S11].

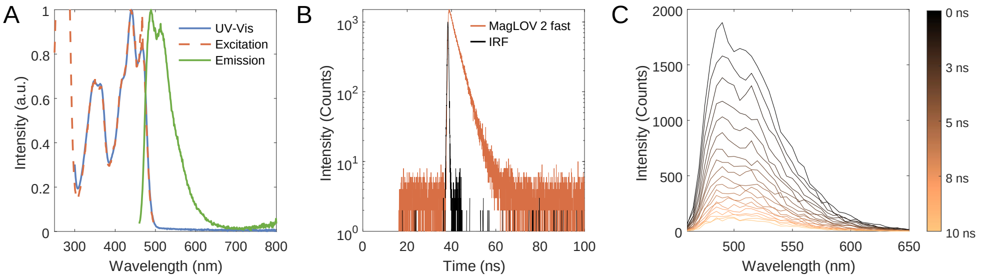


**Fig. S13**: **A**. Normalised UV-Vis absorption spectrum (*blue*), excitation spectrum (em. 510 nm, *orange dash*), and emission spectrum (exc. 450 nm, *green*) for purified solution of MagLOV 2 fast. **B**. Fluorescence decay for MagLOV 2 fast solution (em. 510 nm, *orange*). Instrument response function (IRF) shown in *black*. A straight line on a semilog scale indicates a monoexponential fluorescence decay. **C**. Time-resolved emission spectrum for MagLOV 2 fast solution. Colourbar indicates time after $I_{max}$. TCSPC traces (10 MHz repetition rate) were collected for wavelengths in the range 460-650 nm with 5 nm increments. Global fitting with reconvolution indicates a monoexponential decay with $\tau = 3.245 \pm 0.0002\,\text{ns}$ and a global $\chi^2 = 1.059$.

#### S1.18 Spin Dynamics Simulations

Spin dynamics simulations were performed with the open-source software *RadicalPy*. [37] The MARY simulation, using the function `mary`, incorporated the following Hamiltonian in Liouville space,

$$\hat{\hat{L}} = \hat{\hat{H}}_Z + \hat{\hat{H}}_H + \hat{\hat{K}} + \hat{\hat{R}}, \tag{2}$$

where $\hat{\hat{H}}_Z$ and $\hat{\hat{H}}_H$ are the Zeeman and hyperfine interactions ($\text{FMN}^{\bullet-}$: $^{14}\text{N5} = 0.51$ mT and $\text{WH}^{\bullet+}$: $^{14}\text{N1} = 0.32$ mT), respectively. The chemical kinetics are described by the Haberkorn superoperators $\hat{\hat{K}}$, [38] given by,

$$\hat{\hat{K}} = -\frac{k_S}{2}(\hat{Q}_S \otimes \hat{E} + \hat{E} \otimes \hat{Q}_S) + k_f \hat{E} \otimes \hat{E}, \tag{3}$$

where $k_S$ is the rate for the singlet state of the radical pair to recombine to the ground state ($k_{\text{BET}}$ in Fig. 3 E). $k_f$ is the rate of formation of long-lived radicals from the radical pair ($k_{\text{H/Dep}}$ in Fig. 3 E). $\hat{Q}_S$ and $\hat{E}$ represent the singlet projection operator and the identity matrix, respectively. Relaxation, $\hat{\hat{R}}$, was modelled with two relaxation superoperators which describe two different physical phenomena. The generic "random fields relaxation" (RFR) superoperator, $\hat{\hat{R}}_{\text{RFR}}$, describes relaxation of the three Cartesian components of the electron spin [39] with relaxation rate $k_{\text{RFR}}$,

$$\hat{\hat{R}}_{\mathrm{RFR}} = k_{\mathrm{RFR}}\Big(\frac{3}{2}\hat{E}\otimes\hat{E} - \hat{S}_{1x}\otimes\hat{S}_{1x}^{T} - \hat{S}_{1y}\otimes\hat{S}_{1y}^{T} - \hat{S}_{1z}\otimes\hat{S}_{1z}^{T} - \hat{S}_{2x}\otimes\hat{S}_{2x}^{T} - \hat{S}_{2y}\otimes\hat{S}_{2y}^{T} - \hat{S}_{2z}\otimes\hat{S}_{2z}^{T}\Big). \tag{4}$$

Singlet-triplet dephasing (STD) arises from modulation of the electron-exchange interaction [40]. The relaxation superoperator, with a dephasing rate, $k_{\mathrm{STD}}$, is written as,

$$\hat{\hat{R}}_{\mathrm{STD}} = k_{\mathrm{STD}}\Big(\hat{Q}_S\otimes\hat{Q}_T + \hat{Q}_T\otimes\hat{Q}_S\Big). \tag{5}$$

The two relaxation models influence the MARY and ODMR spectra differently, with increasing rates, where STD primarily broadens the spectral width and RFR reduces the MFE and ODMR contrast percentages.

The rate constant $k_S$, $k_f$, $k_{\mathrm{RFR}}$ and $k_{\mathrm{STD}}$ were tuned to fit the MARY simulations to the experimental data. Subsequently, for ODMR simulation, the kinetic rates $k_S$ and $k_f$ were kept fixed and the relaxation rates $k_{\mathrm{RFR}}$ and $k_{\mathrm{STD}}$ were tuned. Across both simulations, the numerical values of the rates found were consistent with those previously reported on radical pair spin dynamics in flavoproteins [41–44].

The MARY spectrum was reproduced using the Liouville-von Neumann equation,

$$\frac{\partial\hat{\rho}}{\partial t} = -i\hat{\hat{L}}(t)\rho(t), \tag{6}$$

where $\hat{\rho}$ is the density operator. The projection of $\rho(t)$ onto state $S$ is given by the trace of the product of the projection operator, $\hat{Q}_S$, with the density matrix giving the expectation value, $\langle\hat{Q}_S\rangle$, explicitly,

$$\langle\hat{Q}_S\rangle = \mathrm{Tr}\big[\hat{\rho}\,\hat{Q}_S\big]. \tag{7}$$

Where Tr is the trace operation.

Attempts to simulate and fit the experimental data without relaxation were unsuccessful (see Fig. S14). Of the various decoherence/relaxation models used, only the combination of random fields relaxation (RFR) and singlet-triplet dephasing (STD) provided a satisfactory description of the data in the Main text. Fig. S14 shows attempts made to simulate the data without decoherence/relaxation included (purple, MFE $\approx$ 45%, $B_{1/2} \approx 1.5$ mT), including STD, but no RFR (green, MFE $\approx$ 28%, $B_{1/2} \approx 12$ mT), and including RFR, but no STD (blue, MFE $\approx$ 2%, $B_{1/2} \approx 1.7$ mT). Only when relaxation, STD, and RFR were included (as in Fig. S12B) was a good fit achieved (MFE $\approx$ 2%, $B_{1/2} \approx 12$ mT), corresponding to $k_S = 3.6\times10^{6}\ \mathrm{s}^{-1}$, $k_f = 1\times10^{6}\ \mathrm{s}^{-1}$, $k_{\mathrm{RFR}} = 8.2\times10^{6}\ \mathrm{s}^{-1}$, and $k_{\mathrm{STD}} = 1.4\times10^{9}\ \mathrm{s}^{-1}$.

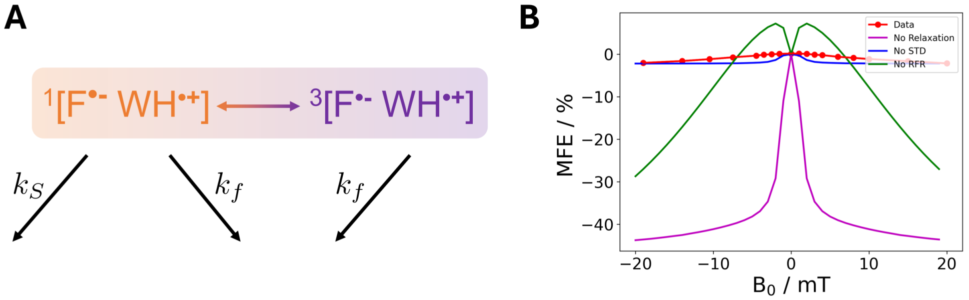


**Fig. S14**: **A.** Reaction scheme used for MARY and ODMR spin dynamics simulations. $k_S$ is the rate for spin-selective recombination to the groundstate. $k_f$ is the rate for the formation of long-lived radicals. **B.** Attempts to simulate and fit the fluorescence-detected MARY spectrum (red) for MagLOV 2 fast using spin dynamics simulations. Purple curve is the simulation with only Haberkorn kinetics included (tuned values of $k_S = 3.6 \times 10^6$ s$^{-1}$, $k_f = 1 \times 10^6$ s$^{-1}$). Blue curve depicts the simulation with Haberkorn kinetics (same as purple) and random fields relaxation (RFR, tuned value of $k_{\text{RFR}} = 8.2 \times 10^6$ s$^{-1}$). The green curve depicts the simulation with Haberkorn kinetics (same as purple) and singlet-triplet dephasing (STD, $k_{\text{STD}} = 1.4 \times 10^9$ s$^{-1}$). Successful simulation was achieved when all four superoperators were included in the simulation (see Fig. S12B).

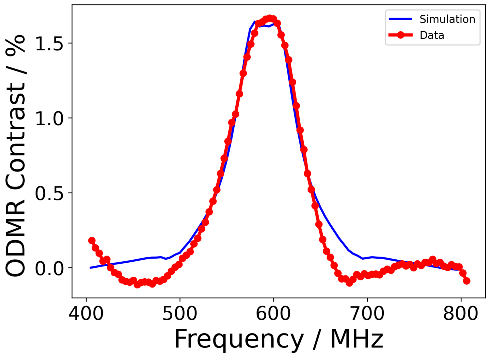


**Fig. S15**: ODMR resonance curve simulation for purified MagLOV 2 fast under 1500 mW/cm$^2$ 450 nm illumination (data, same as Sample 6 in Fig. S6). Kinetic rate parameters $k_S$ and $k_f$ were kept the same as the MARY simulation, while relaxation rates required an adjustment to fit the data, yielding $k_{\text{STD}} = 1.1 \times 10^7$ s$^{-1}$ and $k_{\text{RFR}} = 1.45 \times 10^7$ s$^{-1}$.

For radical pair ODMR simulation [Fig. S15], using the function `odmr` (to be included in a future release of RadicalPy), we employed the rotating frame approximation, which converts a time-dependent Hamiltonian ($\hat{H}(t)$) into a time-independent

Hamiltonian ($\hat{H}_{rot}$).[45] The additional Hamiltonian, $\hat{H}_{rot}$, is as follows,

$$\hat{\hat{H}}_{rot} = \omega_1 \cdot \hat{\mathbf{S}}_{ix} - \omega_{\mathrm{rf}} \cdot \hat{\mathbf{S}}_{iz} - \omega_{\mathrm{rf}} \cdot \sum_j \hat{\mathbf{I}}_{jiz}, \tag{8}$$

where $\omega_1$ is the Larmor frequency of the applied $B_1$ magnetic field; the Coriolis term, $-\omega_{\mathrm{rf}} \cdot \sum_i \hat{\mathbf{L}}_{iz}$, is a fictitious interaction that rationalises the motion of the reference frame. $\hat{\mathbf{L}}$ are the vectors $\hat{\mathbf{S}}$ and $\hat{\mathbf{I}}$ that are the vectors of Cartesian electron and nuclear spin operators, $\{\hat{S}_x, \hat{S}_y, \hat{S}_z\}$ and $\{\hat{I}_x, \hat{I}_y, \hat{I}_z\}$, respectively.

Differences between linewidths in the ODMR and MARY spectra (as noted in Section S1.12) originate from the distinct nature of the two experiments[46]. ODMR directly probes spin resonance transitions between Zeeman-split energy levels of the radical pair. The spectral width is determined by spin relaxation times, inhomogeneous broadening, and microwave field coupling. This often gives a narrower, resonance-defined linewidth. MARY, on the other hand, measures the change in reaction yield or luminescence as a function of an external static magnetic field. In this case, the linewidth is set by the field dependent spin-state mixing and the scale is tied to the hyperfine coupling constants and electron-electron interactions, often giving much broader features. The broader feature reflects the range of fields over which the spin correlation is lost.

To summarise, ODMR predominantly reflects the intrinsic coherence lifetime of the radical pair, leading to slower apparent dephasing and correspondingly narrower spectral features. In contrast, MARY captures ensemble-averaged spin dynamics influenced by additional processes such as recombination and environmental disorder, resulting in faster apparent dephasing and broader magnetic field response curves. These observations are not contradictory, but rather highlight complementary aspects of the same radical pair system. This is reflected in the use of different $k_{\mathrm{STD}}$ values for MARY and ODMR spin dynamics simulations.

#### S1.19 Varying LED parameters

From measurements on earlier variants of MagLOV, we found that increasing the LED power tends to increase the rate at which the MFE approaches saturation (or equivalently, decreases $\tau$). As such, multiplexing experiments and variant characterisation were performed at lower LED power (800 mW/cm$^2$) such that the MFE curves were more distinguishable (i.e. at sufficiently high LED power, different variants begin have converging MFE rates as this effect saturates). Note that this dependence on light source intensity contributes to differences in response times observed in different experimental setups (e.g. microscopy versus bulk measurement instruments) used throughout our work.

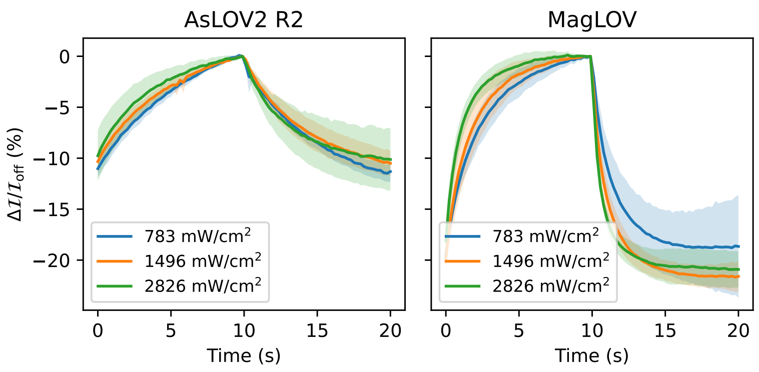


**Fig. S16**: LED characterisation performed on early variants AsLOV2 R2 and MagLOV demonstrate that LED intensity affects MFE saturation rate $\tau$, with higher LED intensities leading to significantly accelerated saturation. Measurements performed on the widefield microscope averaging over single cells as in the main text.

#### S1.20 Spatial Localisation Experiments

The design of the fluorescence MRI instrument used for experiments shown in Fig. 5 C-F are detailed here.

##### S1.20.1 Localisation Optics and Control

The optical path for the MRI setup was constructed using off the shelf parts, custom electronic systems, and 3D-printed components. The same LED and LED driving circuitry are used as in the microscopy experiments, with the LED light at $\sim$10 W. LED light first passes through a 448 nm/20 nm bandpass filter (Edmund Optics, #86-975) and is then coupled into 7 optical fibers (Edmund Optics, #02-536). One of the optical fibers leads to a feedback photodiode for optical stabilisation of light intensity (ensuring excitation illumination is stable within $\sim$ 0.1% to reject heating effects). The remaining 6 fibers carry the light to the sample [Fig. S17 B (ii)]. The sample is a PDMS cast with hollow tubes into which MagLOV cells can be inserted, with the tubes positioned along the $z$-axis [Fig S17 B (i)]. The collection path consists of a quartz rod (1000mm long 10mm diameter, Robson Scientific RQR 10) to allow detection/excitation sensors to be placed far from the sample to eliminate possible interference from RF/Magnetic fields with 475 nm long pass filters (Edmund Optics, #84-737) at each end. The light is then coupled into a USB photodiode (Thorlabs, PM100D). All light conductors (e.g. quartz rod and fibers) were coated in black opaque plastic to avoid external light interference. All couplings and mounts were custom designed and 3D printed in black plastic to block light and (for those near the sample) prevent movement, heating, or other interference as magnetic/RF fields were varied.

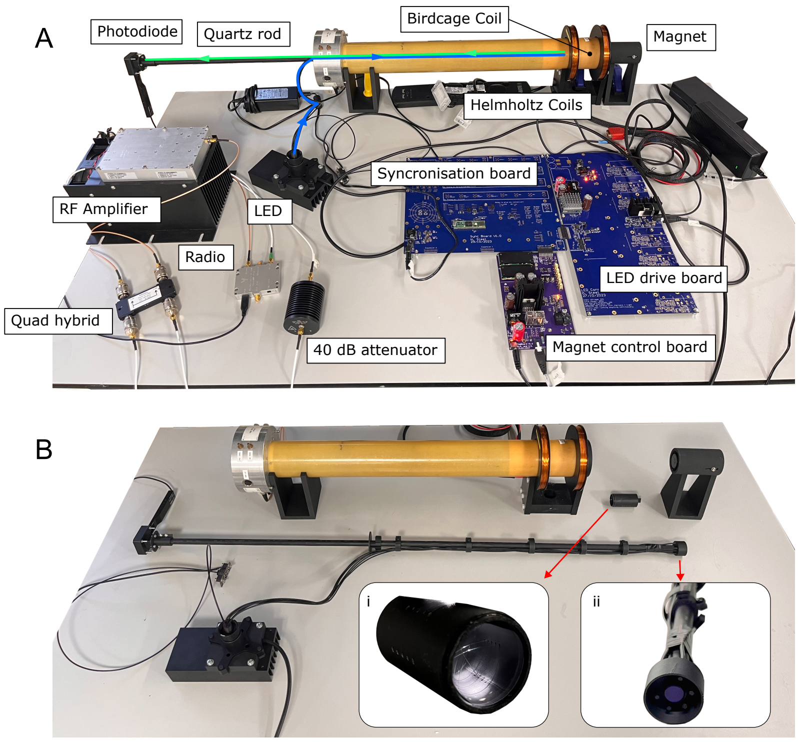


**Fig. S17**: **A, B.** Photographs of the 1D MRI setup, assembled (A.) and disassembled (B.). The LED housing includes a 448 nm/20 nm bandpass between the LED and the fibers. **B.i** The sample holder filled with PDMS, with hollow tubes visible that are loaded with MagLOV cells. **B.ii** The end of the quartz rod at the sample location has 6 fiber optic cables carrying the LED illumination light, and a 475 nm long pass filter on the collection path. Another 475 nm long pass filter is located in the housing attaching the quartz rod to the photodiode.

##### S1.20.2 Localisation RF Hardware

The same RF radio and 20 W amplifier were used for the ESR imaging experiments as the widefield ODMR experiments. This was used to drive a custom-built 28 mm internal diameter quadrature high-bypass “birdcage coil” originally designed for $^1$H imaging at 11.7 T / 500 MHz. This highly resonant structure generates a uniformly circulating $B_1^{\pm}$ field within a cylindrical field-of-view of diameter 28 mm and $z$-extent 40 mm. It is a two-port quadrature design, fed via a high-power RF quadrature hybrid (RADIALL, R432 371) providing RF 90° out of phase. The coil consists of ten cylindrical copper conductors arranged in rungs around a glass-reinforced plastic pullwound tube former with distributed capacitance between them, non-magnetic variable capacitors, and achieves a resonant $Q \approx 30$, and approximately $100\,\mu\text{T}\,\text{W}^{-1}$, and with our 20 W amplifier approximately $B_1 \sim 0.3\,\text{mT}$ was obtained (measured using an $R \approx 1$ mm radius pickup loop). These estimates were obtained by standard

$B_1$ measurement techniques (i.e. fully relaxed multi-angle flip angle calibration) for proton MRI on a separate preclinical 11.7 T MR system, described previously [47].

The neodymium rare earth permanent magnet used for previous ODMR experiments was used to generate an approximately constant linear gradient field measured to be approximately 0.95 T/m, and the isocentre of the measured volume experiment designated at 17.8 mT along this axis. Under these conditions the ESR Larmor frequency is approximately 500 MHz. The RF coil detailed above was used to generate RF at this frequency, and a Helmholtz coil of OD 72 mm was introduced around the outer 68 mm diameter of the RF coil. This Helmholtz coil generates an on-axis $B$-field of approximately $B = \left(\frac{4}{5}\right)^{3/2} \frac{\mu_0 n I}{r}$ where $r = 36\,\mathrm{mm}$, $n = 235$ turns (per coil) and $I$ the delivered current. A constant current was then delivered through the (air-cooled) Helmholtz loop, shifting the resonant condition along the $z$-axis. A DAC and current-controlled amplifier (the same electrical system as described in Fig. S3) was used to provide a varying current $I \in [-1.0, 1.0]$ A.

By varying the Helmholtz coil current between $\pm 1$ A, we can shift the effective magnetic field by $\pm 5.87$ mT around the resonant field of $B_{res} = f/\bar{\gamma}_e = 17.8$ mT. This current modulation brings different spatial positions into resonance while maintaining fixed frequency excitation. A typical range used for imaging was therefore:

$$\Delta x = \frac{2 \times 5.87\,\mathrm{mT}}{0.95\,\mathrm{T\,m^{-1}}} = 12.36\,\mathrm{mm}$$

.

##### S1.20.3 Localisation Data Processing

Imaging was performed in much the same way as for MFE measurements (e.g. Section S1.15.2), however now the RF field is switched on and off rather than the magnetic field (e.g. as was done in Fig. 2 B). A period of 6 s was used, with 6 periods recorded per $B_0$ value, with 28 $B_0$ values scanned. The $B_0$ values were ordered randomly. The data was processed as follows:

1. For each $B_0$ value, a background curve is fit and subtracted, and the periods averaged together yielding $\Delta\mathcal{I}/\mathcal{I}_{\text{off}}$
2. The contrast (i.e. $\Delta\mathcal{I}/\mathcal{I}_{\text{off}}(t = T/2)$) for each $B_0$ point is found
3. A linear trend is subtracted from the contrast values as ordered by their acquisition order (i.e. the corresponding $B_0$ values are in a random order) to account for photobleaching over the course of the experiment
4. The contrast values are then sorted by $B_0$, $B_0$ is converted to $mm$ (as above), and the contrast values are normalised to be in the range $[0, 1]$. This yields the data plot in Fig. 5.
5. The data was then deconvolved with a Gaussian impulse response function of FWHM 80 MHz as previously determined in the widefield ODMR experiments.

Deconvolution is performed using the `richardson_lucy` method in the SciPy package [25].

We note that MRI in general involves a significant amount of signal processing in order to extract the most amount of image information from the signal available [48]. Similarly, to extend fluorescence localisation to three dimensional imaging would require careful signal processing to minimize the number of sampling points required to achieve sufficient spatial resolution before photobleaching degrades the SNR. Achieving this would also motivate future engineering of MagLOV variants with faster rates of contrast change (allowing the off/on RF period to be reduced) and improved photostability.

#### S1.21 Contrast Agent Sensing

Contrast agent experiments were performed on the widefield microscope configured for MFE detection, using microfluidic channels (ibdi $\mu$-Slide VI 0.5 Glass Bottom) to hold samples of Gadobutrol (CRS Y0001803) diluted in PBS buffer in serial dilution (MagLOV concentration the same for all conditions). Measurements were acquired in a randomised sequence (so that photobleaching could be compensated for as in S1.20.3) by programming $6 \times 3$ fields of view (6 imaging conditions, 3 replicates), then automatically performing an MFE measurement acquisition at each field of view. For each MFE acquisition, 10 periods of duration 40 seconds were acquired at 450 nm LED intensity 280 mW/cm$^2$, 100 ms image exposure time and 10 mT magnetic field strength. Optimisation of experimental protocols for for spin-relaxometry based sensing using MagLOV or other radical-pair fluorescent proteins will be required for application, here we demonstrate simply that the spin-radical pair of MagLOV is indeed sensitive to it's surroundings despite the flavin being bound.

#### S1.22 Advantages of Lock-in Detection

Beyond the ability to implement scattering-independent measurements of fluorescent signal *localisation* based on resonance (e.g. as outlined in Section S1.20), MFPs also enable lock-in detection, e.g. modulating a signal-of-interest at a known frequency and then isolating the component of its response at said frequency to reject undesirable noise or background signals (which fall in other parts of the frequency spectrum) to thereby improve sensitivity [49]. Such a method is of great utility for biological assays where a fluorescently tagged molecule or cell is to be distinguished within a tissue or environment that has high signal background [50, 51] - often arising from autofluorescence of the tissue or other neighbouring molecules. Past work investigating this challenge has quantified the practical metric-of-interest for such applications as the Signal-to-Background ratio (SBR). For standard fluorescent labels (e.g. a non-magnetic-responsive fluorescent protein) this can be expressed as $SBR_1 = \frac{n_s}{n_b}$ where $n_s$ is the size of the desired target signal (e.g. in units photons or camera read counts), and $n_b$ is the size of the background signal (e.g. as arising from autofluorescence or other background light sources) [52]. For lock-in based imaging (whether using modulation with a magnetic field or otherwise) we instead have $SBR_2 = \frac{m \times n_s \times \sqrt{N}}{n_b}$ where $m$ is modulation depth defined as the fractional change in $n_s$ when modulation

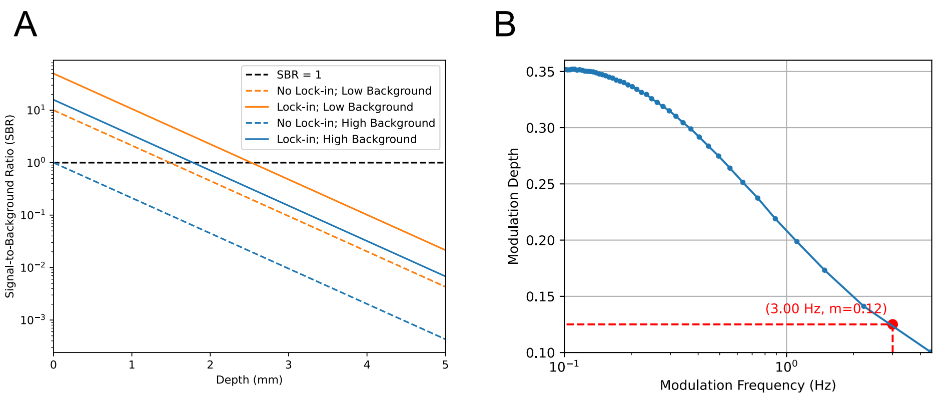


**Fig. S18**: **A.** Analysis of signal-to-background ratio (SBR) with and without lock-in detection in low or high background (i.e. autofluorescent) samples. The black horizontal line indicates SBR = 1, at which the signal of interest is equal to the background. **B.** Plot of simulated modulation depth (i.e. fractional change in fluorescence when magnetic field is applied/removed) of MagLOV 2, based on kinetic model parameters fit to MFE traces (as described in Section S1.15.2).

is applied, and $N$ is the number of samples over which the lock-in measurement is averaged [52]. The form of $SBR_2$ assumes that, while lock-in removes the *constant* component of the background signal $n_b$, its *fluctuations* remain (proportional to square-root of the signal) hence the square-root dependence on $n_b$, and the possibility for improvement in signal quality via averaging more samples (e.g. scaling with $\sqrt{N}$). In contrast $SBR_1$ reflects direct limitation by the size of $n_b$ which isn't possible to "average out" through further data collection. Note that these $SBR$ formulae assume that noise/uncertainty arising from the fluorescence background dominates other noise sources (e.g. shot noise or camera read-out noise) for the purpose of identifying a signal of interest. Further, SBR is distinct from the traditional Signal To Noise (SNR) ratio, in that the minimum SBR for a measurement to be viable can be significantly less than 1 and is highly dependent on the context/system being studied.

To illustrate the advantage of MFE based lock-in detection [49] using MFPs in tissue imaging (for instance, imaging a tagged tumor at depth), Fig. S18 A presents the above metrics for signals arising at different depths in a sample, comparing usage of a standard fluorescent protein against a MFP, and under conditions of low or high (background) autofluorescence. The standard FPs have an intensity $n_{s,0}$ at zero depth of 1000 counts/frametime, whereas the MagLOV FP has an intensity of 500 counts/frametime, reflecting the QE of typical LOV-binding FPs ($\sim 0.2 - 0.4$) being roughly half that of GFP (0.6 and above) [53]. The measured (signal) fluorescence is then $n_s = n_{s,0}e^{-A}$ where $A$ is the total absorbance on the excitation and emission paths. We have $A = A(\lambda_{exc.}) + A(\lambda_{em.})$ where $\lambda_{exc.} = 450$ nm is the excitation wavelength $\lambda_{em.} = 500$ nm is the emission wavelength. Using the data reported in [54] for a rabbit cortex, we have $A(\lambda) = \mu_A(\lambda) \times d$ where $\mu_A \approx 8$ cm$^{-1}$ for excitation and $\mu_A \approx 7.5$ cm$^{-1}$ for emission and $d$ is depth in cm. In the low background case, we estimate the autofluorescence (which arises from the sample as a whole and hence is not scaled with depth) at 10% of the signal fluorescence, i.e. $n_b = 100$ counts/frametime, and in the high background case we estimate the autofluorescence at 100% of the signal fluorescence, i.e. $n_b = 1000$ counts/frametime. We use a modulation frequency of 3 Hz for which MagLOV 2 would exhibit a $\sim 12\%$ modulation depth (hence $m = 0.12$) as in Fig. S18 B, and assume a frametime of 100 ms, and total integration time of 10 s. The results show that, in the low background case, the no lock-in measurement has a $SBR \approx 1$ at approximately 1 mm depth - similar to depths at which past studies have shown ability to localise fluorescent markers in tissue [55]; in our simulation use of lock-in approximately doubles this. In the high background case, lock-in becomes significantly more beneficial, moving the point where $SBR \approx 1$ from the surface to depths of 2 mm. A target of $SBR \approx 1$ may in practice be highly conservative - for example lock-in imaging approaches can function even if a signal-of-interest is many orders of magnitude weaker than background contributors to the measurement (i.e. $SBR \ll 1$) - for example past work has demonstrated this in applications where a signal of interest is less than 0.1% of the total signal [50]. Furthermore, these methods could be improved through greater averaging, or engineering of faster responding/greater modulation depth MFPs; the present analyses are intended only to show how MFPs offer a easy-to-implement method for realising lock-in detection to improve practical imaging performance as explored in past studies [52].

#### S1.23 Lock-In Classification Data

Summary data for Main Text Fig. 4 G is shown in Tables S4 and S5. Balanced accuracy is calculated according to the formula,

$$\text{Balanced Accuracy} = \frac{1}{2}\left(\frac{TP}{TP + FN} + \frac{TN}{TN + FP}\right) \tag{9}$$

where $TP$ = True Positive (red and lock-in), $TN$ = True Negative (not red and no lock-in), $FP$ = False Positive (not red and lock-in) and $FN$ = False Negative (red and no lock-in).

| | True | False |
|---|---|---|
| Positive | 109 | 15 |
| Negative | 1275 | 2 |

**Table S4**: Data for Main Text Fig. 4 G, By Cells

| | True | False |
|---|---|---|
| Positive | 28 | 3 |
| Negative | 201 | 0 |

**Table S5**: Data for Main Text Fig. 4 G, By Trenches

#### S1.24 MFP Variants Engineered by Directed Evolution

| **Variant** | **Mutations** | |
|---|---|---|
| | **Position relative to ancestor** | **Position relative to new protein** |
| AsLOV2 WT | - | - |
| AsLOV2 C450A | $C_{450}A$ | $C_{48}A$ |
| AsLOV2 R1 | $C_{450}A$ $D_{540}M$ | $C_{48}A$ $D_{138}M$ |
| **AsLOV2 R2** | $C_{450}A$ $D_{540}M$ $Q_{513}A$ | $C_{48}A$ $D_{138}M$ $Q_{111}A$ |
| AsLOV2 R3 | $V_{416}T$ $C_{450}A$ $D_{540}M$ $Q_{513}A$ $L_{496}V$ | $C_{48}A$ $D_{138}M$ $Q_{111}A$ $L_{94}V$ |
| AsLOV2 R4 | $C_{450}P$ $D_{540}M$ $Q_{513}A$ $L_{496}V$ | $C_{48}P$ $D_{138}M$ $Q_{111}A$ $L_{94}V$ |
| AsLOV2 R5 | $C_{450}P$ $D_{540}M$ $Q_{513}K$ $L_{496}V$ | $C_{48}P$ $D_{138}M$ $Q_{111}K$ $L_{94}V$ |
| **MagLOV** (R6) | $C_{450}P$ $D_{540}M$ $Q_{513}K$ $L_{496}V$ $G_{528}K$ | $C_{48}P$ $D_{138}M$ $Q_{111}K$ $L_{94}V$ $G_{126}K$ |
| MagLOV R7 | $C_{450}P$ $D_{540}M$ $Q_{513}K$ $L_{496}V$ $G_{528}K$ $R_{448}W$ | $C_{48}P$ $D_{138}M$ $Q_{111}K$ $L_{94}V$ $G_{126}K$ $R_{46}W$ |
| MagLOV R8 | $C_{450}P$ $D_{540}M$ $Q_{513}K$ $L_{496}V$ $G_{528}R$ $R_{448}W$ | $C_{48}P$ $D_{138}M$ $Q_{111}K$ $L_{94}V$ $G_{126}R$ $R_{46}W$ |
| **MagLOV 2** (R10) | $C_{450}P$ $D_{540}M$ $Q_{513}K$ $L_{496}V$ $G_{528}R$ $R_{448}W$ $E_{525}G$ $F_{494}L$ | $C_{48}P$ $D_{138}M$ $Q_{111}K$ $L_{94}V$ $G_{126}R$ $R_{46}W$ $E_{123}G$ $F_{92}L$ |
| **MagLOV 2 fast** | $C_{450}P$ $D_{540}M$ $Q_{513}K$ $L_{496}V$ $G_{528}R$ $R_{448}W$ $E_{525}G$ $F_{494}L$ $A_{543}L$ | $C_{48}P$ $D_{138}M$ $Q_{111}K$ $L_{94}V$ $G_{126}R$ $R_{46}W$ $E_{123}G$ $F_{92}L$ $A_{141}L$ |

**Table S6**: Comparison of mutations relative to ancestor protein (AsLOV2 $C_{450}A$ [2]) sequence, as well as referenced to beginning of MagLOV stand-alone variant (which removes the first 402 residues). Variants in **bold** are those characterised in this paper (e.g. main text Fig. 2); AsLOV2 R5 is also used in main text Fig. 4 B,C.
